# SRSF1 as a novel interacting partner for IFITM1/3 unravels the emergent role of IFITM1/3 mediating protein expression

**DOI:** 10.1101/2022.02.02.478792

**Authors:** Maria Gómez-Herranz, Jakub Faktor, Marcos Yébenes Mayordomo, Magdalena Pilch, Lenka Hernychova, Kathryn L. Ball, Borivoj Vojtesek, Ted R. Hupp, Sachin Kote

## Abstract

IFITM proteins play a role in cancer cell progression through undefined mechanisms. Here, we propose an emergent role of IFITM1/3 regulating protein synthesis. SBP-tagged IFITM1 protein was used to identify an association of IFITM1 protein with the cytosolic isoform of SRSF1 that transports mRNA to the ribosome. This cytosolic association was confirmed in situ using proximity ligation assays for SRSF1 and IFITM1/3, suggesting a role associated with translation. Accordingly, IFITM1/3 were shown to interact with HLA-B mRNA in response to IFNγ stimulation using RNA-protein proximity ligation assays. In addition, shotgun RNA sequencing in IFITM1/IFITM3 null cells and wt-SiHa cells indicated that reduced HLA-B gene expression does not account for lowered HLA-B protein synthesis in response to IFNγ. Furthermore, ribosome profiling using sucrose gradient sedimentation identified a reduction in 80S ribosomal fraction an IFITM1/IFITM3 null cells compared to their wild-type counterpart, partially reverted by IFITM1/3 complementation. Our data all together link the binding of IFITM1/3 proteins to HLA-B mRNA and SRSF1 as a mechanism to catalyze the synthesis of target proteins, suggesting an RNA chaperonin role for IFITM1/3 proteins.

**Significance:** IFITMs are widely studied for their role in inhibiting viruses, and multiple studies have associated IFITMs with cancer progression. However, mechanistic insights are not well understood. Our study proposes that IFITMs have a role regulating protein synthesis, a pivotal function highly relevant for viral infection and cancer progression. Our results suggest that IFITM1/3 is present in the ribosomal fraction and regulates particular protein expression; among them, we identified HLA-B. Changes in HLA-B expression could impact the presentation and recognition of oncogenic antigens on the cell surface by cytotoxic T cells and, ultimately, limit tumor cell eradication. In addition, the role of IFITMs mediating protein translation is relevant, as has the potential of regulating the expression of viral and oncogenic proteins.

## Introduction

Multiple oncogenic pathways selectively modulate the expression of mRNAs, and in addition, regulation of protein expression is essential for cancer development (Robichaud *et al.*, 2019). Thus, there is a particular scientific interest to unravel which cellular components orchestrate the translation of specific mRNA products associated with tumor progression. Indeed, control of cancer-specific translation is emerging as a new anticancer strategy where therapeutic drugs inhibit mRNA translation in a selective manner (Xu and Ruggero, 2020).

Furthermore, interferons (IFNs) form a family of cytokines originally discovered to respond in opposition to the adverse effects of the flu virus; however, subsequent biological roles described for IFNs include activity against cancer development (Platanias, 2005). In this regard, IFNs are used in cancer treatment (Tamura *et al.*, 1987; Pujade-Lauraine *et al.*, 1996; Windbichler *et al.*, 2000; Marth *et al.*, 2006; Alberts *et al.*, 2008; Parker, Rautela and Hertzog, 2016; Ives *et al.*, 2017; van der Kooij *et al.*, 2020) due to their ability to inhibit proliferative pathways, promote cellular apoptosis and increase the activity of the immune system (Budhwani, Mazzieri and Dolcetti, 2018).

Although relatively elevated doses of IFNs show protective effects against tumor progression (Wang, Rahbar and Fish, 2011; Booy, Hofland and van Eijck, 2015; Green *et al.*, 2016), continuous low production of IFN stimulates the expression of the interferon resistance DNA damage signature (IRDS). The IRDS is comprised of a small subset of ISGs (Weichselbaum *et al.*, 2008; Cheon, Borden and Stark, 2014) that promote tumor development by acquiring resistance to DNA damage, scaping immune surveillance, and increasing metastatic spread (Wallace, Martin and Ambs, 2011; Kolosenko *et al.*, 2015). Thus, IFNs can mediate both cancer suppression and growth depending on context (Jorgovanovic *et al.*, 2020).

There are described three protein families within the mammalian IFN tree; type I (IFNα, IFNβ, IFNe, IFNk and IFNω), type II (IFNγ) and type III (IFNλ) (Bekisz *et al.*, 2004; Borden *et al.*, 2007). The core mechanism of action of IFNs involves activating the JAK kinase-STAT signaling pathway that induces the transcription of numerous interferon-stimulated genes (ISGs) (Williams, 1991; Darnell, Kerr and Stark, 1994), including IFITM1/2/3.

The immune-related interferon-induced transmembrane (IFITM) protein family are composed of three members; IFITM1, IFITM2, and IFITM3 (Bailey *et al.*, 2014). IFITM1 is slightly different from IFITM2 and IFITM3, with some studies proposing that IFITM1 is uniquely expressed on the cell surface (Weston *et al.*, 2014; Jia *et al.*, 2015). In addition, only a limited number of interacting partners have been identified for the IFITMs (Xu, Yang and Hu, 2009; Amini-Bavil-Olyaee *et al.*, 2013; Narayana *et al.*, 2015; Yu, Xie, Ng, Lum, M.-Y. Cai, *et al.*, 2015). However, a growing body of research has proven that components of the IFITM family are capable of attenuating the propagation of RNA virus such as influenza A virus (IAV), West Nile virus (WNV), dengue virus, Severe Acute Respiratory Syndrome (SARS) coronavirus, filoviruses, Vesicular Stomatitis Virus (VSV), and Hepatitis C virus (HCV) (Brass *et al.*, 2009; Bailey *et al.*, 2014).

In addition to the widely studied anti-viral function of the IFITMs, IFITM1 and IFITM3 also function as a pro-oncogenic proteins whose expression has been reported in various cancers such as breast, cervix, colon, leukemia, ovary, brain, and esophagus (Fan *et al.*, 2008; Györffy *et al.*, 2008a; Weichselbaum *et al.*, 2008; Wu *et al.*, 2011; Borg *et al.*, 2016a; Ogony *et al.*, 2016; Sari, Y.-G. Yang, *et al.*, 2016; Liu *et al.*, 2020; Wang *et al.*, 2020). Moreover, IFITM1 is an IRDS gene expressed upon radiation resistance. Hence, the protective effect described for IFNs against tumors is not well reflected in the role of the IFN-induced IFITM proteins concerning cancer (Weichselbaum *et al.*, 2008; Khodarev, Roizman and Weichselbaum, 2012). Nonetheless, the molecular mechanism whereby IFITM1 and IFITM3 promote cancer growth or mediates viral restriction is not well-defined.

Overall, the IFITM functions associated with blocking viral infection, IFITM proteins can reduce HIV-1 viral protein synthesis by preferentially excluding viral mRNA transcripts from translation (Lee *et al.*, 2018) providing an intracellular function for the IFITM family linked to suppressed viral propagation. This is consistent with our previous research where we found a translational role for the IFITM family in the context of cancer; deletion of IFITM1/3 genes suppresses the expression of a subset of proteins whose synthesis is mediated by IFNγ. Interestingly, HLA-B, which is a component of the IRDS was significantly identified (Gómez-Herranz *et al.*, 2019). This indicates that IFITM1/3 can regulate the synthesis of some anti-viral in addition to cancer-associated gene products.

In this report, we further define the molecular mechanism whereby genetic ablation of IFITM1 and IFITM3 attenuates the synthesis of a subset of IFN-responsive proteins despite IFITM1/3-knockout cell still retaining the ability to mediate IFN-induced gene expression. Here, we show that IFITM1/3 proteins can interact with the cytosolic translation factor SRSF1 and HLA-B mRNA. In addition, we identified changes in the 80S ribosomal fraction in the absence of IFITM1/3 expression. Taking all together, this provides additional support for an RNA-binding and translational role for the IFITM family of proteins, as reported in response to HIV infection (Lee *et al.*, 2018).

## Materials and Methods

### Cell culture

The wt-SiHa cells and IFITM1/IFITM3 null cells, that were described previously (Gómez-Herranz *et al.*, 2019) were grown in RPMI 1640 medium (Invitrogen, USA) supplemented with 10% (v/v) fetal bovine serum (Labtech, UK) and 1% penicillin/streptomycin (Invitrogen, USA) and incubated at 37 °C with 5% CO_2_.

### Western blotting

Protein from lysed samples was quantified using the Bradford (Bradford, 1976) Protein Assay Dye Reagent (Bio-Rad, USA). Proteins were resolved by SDS-PAGE using 15% gels and transferred onto nitrocellulose membranes (Amersham Protran, GE Healthcare, USA). Immunoblots were processed by enhanced chemiluminescence (ECL) and quantified as RLU.

### Antibodies

Proteins were detected using the following primary antibodies: mouse monoclonal antibodies were generated to a peptide that is identical in IFITM1 and IFITM3 and their characterization was described previously (Gómez-Herranz *et al.*, 2019). As this panel of monoclonal antibody cannot distinguish between the IFITM1 and IFITM3 proteins, the text specifically states that IFITM1 and IFITM3 proteins (shortened to IFITM1/3) were measured when using these antibodies. Other sources of antibodies include Mouse monoclonal anti-IFITM2 (Proteintech), rabbit polyclonal anti-SRSF1 (Thermo Fisher Scientific), rabbit polyclonal anti-HLA-B (Thermo Fisher Scientific), rabbit polyclonal anti-RPL7a (Cell Signaling), mouse monoclonal anti-β-ACTIN (Sigma-Aldrich) and mouse monoclonal anti-GAPDH (Abcam).

### Trichloroacetic acid (TCA) precipitation

Proteins were precipitated from individual sucrose gradient fractions by TCA precipitation. While adding 10 μl of 1 μg/μl of Bovine Serum Albumin (BSA) to each prechilled eluate followed by the addition of ice-cold TCA (90 μl), then was mixed and incubated on ice for 60 min. The fractions were then centrifuged at 10,000 g at 4ºC for 10 min. The pellet was again centrifuged to remove any residual supernatant, washed with 250 μl of ice-cold acetone and centrifuged at 4ºC for 5 min. Finally, the supernatant was discarded and a visible small white precipitate was air dried. Protein was then resuspended in 2x sample buffer to proceed with the immunoblot.

### Immunofluorescence

The wt-SiHa and IFITM1/IFITM3 null cells were non-stimulated and stimulated with 100 ng/ml IFNγ for 24 h. Cells were fixed with 4% (v/v) paraformaldehyde in PBS at RT for 15 min, washed with PBS three times, and permeabilized using 0.25% triton X-100 in PBS at RT for 10 min. Then, the cells were again washed with PBS three times and blocked with 3% BSA in PBS for 1 h. The primary antibody was incubated at appropriate dilution (typically 1:1000) overnight at 4 ºC. Depending on which host species the primary antibody been generated, Alexa Fluor 488 goat anti-mouse (Invitrogen). Alexa Fluor 594 donkey anti-rabbit (Invitrogen) or Alexa Fluor 594 goat anti-mouse (Invitrogen) secondary antibody was incubated at RT for 1 h. Coverslips were washed three times with PBS in between each step. Cells were incubated in DAPI (Invitrogen) diluted at 1:10,000 with dH_2_O for 5 min to stain the nucleus. An additional three washes with dH_2_O for 5 min were performed. A single drop of Fluorescence Mounting Medium (S3023, Dako, Denmark) was used to mount the cells on the slide. The fluorescent signal was detected using a Zeiss Axioplan 2 microscope (63x or 100x oil immersion objective). Images were acquired by Micro-Manager 1.4 software. Images were processed in ImageJ 2.0 software.

### RT-qPCR

The following parameters were considered while designing the primers: Firstly, primers were designed to the codifying region of IFITM1 (F: 5’-ACTGGTATTCGGCTCTGTGAC-3’; R: 5’-GCTGTATCTAGGGGCAGGAC-3’), IFITM3 (F: 5’-CAAACCTTCTTCTCTCCTGTCAA-3’; R: 5’-GATGTGGATCACGGTGGAC-3’), IRF1 (F: 5’-CTCTGAAGCTACAACAGATGAG-3’; R: 5’-GTAGACTCAGCCCAATATCCC-3’), ISG15 (F: 5’-GAGGCAGCGAACTCATCTTT-3’; R: 5’-AGCATCTTCACCGTCAGGTC-3’), HLA-B (F: 5’-CACTGAGCTTGTGGAGACCA-3’; R: 5’-ATGACCACAACTGCTAGGACA-3’), B2M (F: 5’-CTCGCTCCGTGGCCTTAG-3’; R: 5’-GGATGAAACCCAGACACATAGC-3’), STAT1 (F: 5’-CCATCCTTTGGTACAACATGC-3’; R: 5’-TGCACATGGTGGAGTCAGG-3’), and β-ACTIN (F: 5’-CATGTACGTTGCTATCCAGGC-3’; R: 5’-CTCCTTAATGTCACGCACGAT-3’) genes. Designing IFITM1 and IFITM3 primers target the unique C-terminal and N-terminal regions respectively; the specific gene products were as follows, including gene name, gene number, transcript, transcript number; IFITM1; ENSG00000185885, IFITM1-202, ENST00000408968.4; IFITM3, ENSG00000142089, IFITM3-201, ENST00000399808.5; IRF1, ENSG00000125347, IRF1-201, ENST00000245414.9; ISG15, ENSG00000187608; ISG15-201, ENST00000379389.4; HLA-B, ENSG00000234745, HLA-B-249, ENST00000412585.7; B2M, ENSG00000166710, B2M-204, ENST00000558401.6; STAT1, ENSG00000115415, STAT1-201, ENST00000361099.7; β-ACTIN, ENSG00000075624, ACTB-201, ENST00000331789.11.

Additional parameters were that all amplicons were expected to have a product size around 150–250 bp and with a melting temperature difference under 1ºC between primers, as multiple genes were run in parallel using the same PCR plates. Designed primer sequences were analyzed by BLAST to ensure there were no additional amplified products. RT-qPCR was run under the same conditions for all the different transcripts. Melting curve analysis was performed, to define the specificity of each primer pair (data not shown). Three technical replicates from three biological replicates were set for each sample condition and the raw Ct values of all samples were defined (data not shown). To allow a stringent analysis, Ct expression values were less than one unit difference between each technical triplicate. Relative Ct expression values were normalized with a housekeeping gene, b-ACTIN. The analysis of gene expression was taken from the Ct values obtained by RT-qPCR, normalized into a relative expression using the formula 2^-DCt^ where DCt is the difference between the Ct of the transcript of interest and the Ct of b -ACTIN (Fig 3A).

### Proximity ligation assay (PLA)

The wt-SiHa and IFITM1/IFITM3 null cells were grown and processed as described in the immunofluorescence method. Primary antibodies pairs from different species were incubated overnight on the fixed, permeabilized, and blocked cells. Depending on requirements of the experiment, the following combination of antibodies were incubated for 18 h at 4ºC: the IFITM1/IFITM3 mouse monoclonal (1:500 dilution) with the SRSF1 rabbit polyclonal (1:250 dilution), or the IFITM1/IFITM3 mouse monoclonal (1:500 dilution) with the rabbit monoclonal anti-BIOTIN (1:200 dilution); the IFITM1/IFITM3 mouse monoclonal (1:500 dilution) with the rabbit polyclonal anti-RPL7a (1:250 dilution). Samples were incubated with the PLA probes; probe-mouse MINUS (DUO92004, Sigma-Aldrich, USA) and probe anti-rabbit PLUS (DUO92002, Sigma-Aldrich, USA) at 37ºC for 1 h. Samples were washed three times with buffer A (150 mm NaCl, 10 mM Tris Base, 0.05% (v/v) Tween-20, pH 7.4) for 5 min and then samples were incubated with ligation buffer (8 μl of 5x ligation stock (New England Biolabs, USA), 1 μl ligase and 31 μl of ultrapure water on each coverslip) at 37 ºC for 30 min. Samples were again washed three times with buffer A for 5 min and incubated with amplification buffer at 37ºC for 2 h. Reagents are from Duolink in Situ Detection Reagents Green assays (DUO92014, Sigma-ALdrich, USA). Samples were washed twice with buffer A, three times with buffer B (100 mM NaCl, 50 mM Tris base, pH 7.5), and once with 0.01x buffer B. Cells were incubated in DAPI (Invitrogen) diluted at 1:10,000 with 0.01x buffer B for 5 min to stain the nucleus. An additional 2 washes with 0.01x buffer B for 5 min were performed prior to mounting. The fluorescent signal was detected using a Zeiss Axioplan 2 microscope (63x or 100x oil immersion objective). Images were acquired by Micro-Manager 1.4 software. Images were processed in ImageJ 2.0 software.

### RNA in situ hybridization-PLA (rISH-PLA)

The wt-SiHa cells and IFITM1/IFITM3 null cells were non-stimulated and IFNγ-stimulated for 24 h. Cells were fixed with 4% paraformaldehyde in PBS at RT for 20 min and washed with PBS for 10 min. Then, samples were incubated in 70% (v/v) ethanol overnight. Cells were then washed in PBS for 30 min and permeabilized with 0.05% CHAPS and 0.4% Triton X-100 for 10 min at RT. Next, samples were treated with hybridization buffer for 30 min at RT. Samples were incubated for hybridization with 40 μl of hybridization buffer (10% (v/v) formamide, 2X SSC, 0.2 mg/mL E. coli 522 tRNAs, 0.2 mg/mL sheared salmon sperm DNA and 2 mg/mL BSA) containing 50 ng of or HLA-B-biotin DNA probe (5’ TGTCCTAGCAGTTGTGGTCATCGGAGCTGTGGTCGCTGCTGTGAT-biotin 3’) (Sigma-Aldrich, USA) in a humidified chamber overnight. Prior to that, 5 μl of probe diluted in water was denatured for 5 min and chilled on ice for 5 min. Then, samples were washed with 2X SSC and 10% formamide at RT, twice with hybridization buffer at 37 ºC, 2X SSC at RT, and finally with PBS at RT. Each wash was carried out for 20 min. Next, samples were blocked with 3% BSA and 0.1% saponine in PBS at RT for 30 min. After that, they were incubated with rabbit anti-BIOTIN and mouse-anti-IFITM1/IFITM3 at RT for 2 h. The subsequent steps were performed as described in the PLA methodology section (above).

### Bioinformatic analysis of the RNA-sequencing

Biological triplicates of wt-SiHa cells and IFITM1/IFITM3 null cells were non-stimulated and stimulated 24 h with 100 ng/ml IFNγ. Total RNA was extracted from frozen cell pellets following the instruction manual (RNeasy Mini kit, Qiagen, Germany). RNA samples of IFNγ-stimulated and non-stimulated wt-SiHa cells and IFITM1/IFITM3 null cells were processed by Otogenetics (USA, Georgia) for paired end RNA sequencing analysis, using an Illumina HiSeq 2500 and designated 20 million reads. The pair of fastq files obtained from the sequencer were checked for quality control, merged and processed using CLC Genomic Workbench 12.0 to obtain the total RNA expression levels. GRCh38 was taken as the human reference genome with the following settings: mismatch cost: 2, insertion cost: 3, deletion cost: 3. The results were compared using RNA gene expression for each condition, creating four different scatter plots (Fig 3B) that compared IFNγ stimulated to nonstimulated conditions, and wt-SiHa cells to IFITM1/IFITM3 null cells. Comparisons were performed taking the transcripts per million (TPM) values as reporting abundances and Log2 (TPM condition 1 versus Log2 (TPM condition 2) as comparison values. An additional heat map was generated with the list of IRDS genes to compare the gene expression of these genes of interest across samples (SI Appendix, Fig. S4). Following the same parameters, another heat map was created using ggplot2 to compare isoform switches using sum of RNA transcript expression or the RNA transcript expression level of particular transcripts. The color code represented in the heat map shows; red when a gene is highly expressed and purple to blue for non-expressed and under-expressed values. The values in white represent a low level of expression becoming non-significant. Finally, the values in grey are for the genes in which TPM was equal to 0.

### Sucrose gradient sedimentation

For experiments processing ribosomal fractions using the wt-SiHa cells and IFITM1/IFITM3 null cells (Fig 5), the cultures were stimulated with 100 ng/ml IFNγ for 24 h prior to cell harvesting. In the complementation assays (SI Appendix, Fig. S5D-F), whereby IFITM1/IFITM3 null were transfected with either the IFITM1 and IFITM3 expression plasmids or empty vector controls, 24 h after transfection, the cells were then stimulated with 100 ng/ml IFNγ for 24 h. In all cases, the cells were treated with 50 μg/ml cycloheximide (Merck Chemicals, Germany) for 30 min. Then, cells were washed in PBS (phosphate buffered saline; 137 mm NaCl, 2.7 mM KCL, 10 mm Na2HPO4, and 1.8 mm KH2PO4) containing 1x RSB, harvested by centrifugation at 7,000 rpm for 1 min at 4 °C, and frozen at −80 ºC. Lysis was carried out by resuspending the cell pellets in 250 μl of RSB/RNasin buffer and 250 μl PEB (Polysome extraction buffer) buffer. Mechanical disruption of the cell lysate was carried out by passing the lysate though a needle (25G) five times. Lysates were incubated on ice for 10 min and centrifuged at 10,000 g for 10 min at 4ºC. Lysates were processed as described (Sanford *et al.*, 2004). Sucrose gradients (10-45%) were prepared using a BioComp gradient master. Lysates were applied onto the gradient and centrifuged at 41,000 rpm for 2 h 30 min using a SW41 rotor. Fraction collection was performed using a BioComp gradient station model 153 (BioComp Instruments, Canada). Analysis of ribosomal fractions was performed as biological triplicate. The 10X RSB stock solution contained 200 mM Tris-Hcl (pH 7.5), 1 M KCL, and 100 mM MgCl2. The RSB/RNasin buffer (2 ml) was prepared fresh and contained 200 ul of 10 X RSB and 50 ul RNasin (N2511, Promega, USA). The PEB buffer (10 ml) was made fresh and contained 1 ml of 10 x RSB, 50 ul of NP40, and 1 protease inhibitor tablet (Complete Mini-EDTA free EASYpack, 034693159001, Roche/Sigma-Aldrich). In addition, sample preparation for western blotting required deproteinization of crude samples and was performed using the trichloroacetic acid (TCA) method.

### Cloning, transfection and affinity purification of IFITM1 from heavy isotope labeled cells

IFITM1 cDNA was cloned by PCR into pEXPR-IBA105 expression vector containing a SBP tag at the N-terminus of the coding region (SBP vector, IBA, Germany). Cells were grown as biological triplicates for 10 days with 5 passages in RPMI SILAC media before transfection (Dundee Cell Products, UK). Cells were isotopically labelled with light media; L-[12C614N4] arginine (R0) and L- [12C614N2] lysine (K0) and heavy media; L-[13C614N4] arginine (R6) and L-[13C614N2] lysine (K6). For transfection, cells were grown to approximately 80% confluency in light and heavy media and transfected using Attractene (#301007, Qiagen, Germany) with SBP-empty vector (control cells) and SBP-IFITM1 (SI Appendix, Fig. S2). At 24 and 48 h after transfection, cells were washed twice in ice cold PBS and scraped into 0.1% Triton buffer for 30 min on ice. Equal amounts of protein were used for performing the pull down. Total protein extracts were measured by Bradford assay. For affinity purification, the cells were washed twice in cold PBS and scraped directly into IP buffer (100 mM KCl, 20 mM HEPES pH 7.5, 1 mM EDTA, 1 mM EGTA, 0.5 mM Na3VO4, 10 mM NaF, 10% (v/v) glycerol, protease inhibitor mix, and 0.1% Triton X-100). The lysate was then incubated for 30 min on ice and centrifuged at 13,000 rpm for 15 min at 4 °C. Then cell lysate was added to Streptavidin Agarose conjugated beads (Millipore, USA) and incubated for 2 h with gentle rotation. The proteins were eluted from the beads using Elution Buffer (20 mM HEPES pH 8, 2 mM DTT, and 8 M Urea). The eluted samples (whole volume) from light and heavy SILAC medium labeled cells were mixed together.

### Peptide generation using FASP

Proteins eluted after SBP pull-down were processed by the filter-aided sample preparation protocol method (FASP) (Wiśniewski *et al.*, 2009) according to a workflow described in Gómez-Herranz et al. (Gómez-Herranz *et al.*, 2019). Briefly, protein concentration was determined using the RC-DC protein assay (Bio-rad, USA). Approximately 100 μg of protein dissolved in 20 mM HEPES pH 8, 2 mM DTT, and 8 M Urea was added to a 10 kDa spin filter column (Microcon, Merck-Millipore, USA), on-filter reduced, alkylated and trypsin digested to peptides. Tryptic peptides were then desalted using C18 micro-spin columns (Harvard Apparatus, USA) (Gómez-Herranz *et al.*, 2019).

### LC-MS/MS analysis of SILAC labeled samples

Tryptic peptides from isotopically labeled cells were separated using an UltiMate 3000 RSLCnano chromatograph (Thermo Fisher Scientific, USA). Tryptic peptides were loaded onto a pre-column (μ-precolumn, 30 μm i.d., 5 mm length, C18 PepMap 100, 5 μm particle size, 100 Å pore size) and separated using an Acclaim PepMap RSLC column (75 μm i.d., length 500 mm, C18, particle size 2 μm, pore size 100 Å). Tryptic peptides were separated by a linear gradient of mobile phase B (B = 80% *(v/v)* acetonitrile (ACN), 0.08% *(v/v)* formic acid (FA) in water) and A (A = 0.1% *(v/v)* FA in water) as follows: 2 % B over 4 min, 2 – 40 % B over 64 min, 40 – 98 % B over 2 min. The flow rate was 300 nl/min. Tryptic peptides eluting from the column were injected into an Orbitrap Elite (Thermo Fisher Scientific, USA) operating in Top20 data-dependent acquisition mode. Scanning was set to 400 – 2000 m/z performed at 120 000 resolution. The AGC target was 1 x 10^6^ with a 200 ms injection time and twenty data-dependent MS2 scans (1 microscan, 10 ms injection time and 10 000 AGC).

### Database searching and analysis

The SILAC data were processed using Proteome Discoverer 1.4 (Thermo Fisher Scientific, USA) and the Mascot search engine with the following search settings: a human database, Swiss-Prot (April 2016); enzyme-trypsin; 2 missed cleavage sites; a precursor mass tolerance of 10 ppm; a fragment mass tolerance of 0.6 Da; modification included: carbamidomethyl [C], oxidation [M], acetyl [protein N-terminus]. The search results were used to generate the final report with a 1% FDR on both PSM and peptide groups. SILAC labels of R6 and K6 were chosen for heavy and R0 and K0 for light. The relative quantification value was represented as heavy/light ratio (Table S1).

### Data availability

The mass spectrometry proteomics data have been deposited to the ProteomeXchange Consortium via the PRIDE partner repository with the dataset identifier PXD030540, Username, Password: jM0wyPd7. The entire set of RNA seq files have been deposited to the datadryad.org with DOI https://doi.org/10.5061/dryad.c59zw3r92.

## Results

### Developing an isogenic cervical cell model

IFITMs are IFN-stimulated molecules involved in human cancer development, including cervical carcinogenesis (Pan *et al.*, 2010). In addition, gene expression of cervical cancer specimens reveals an inverse correlation between IFNγ and lymph node metastases (Kim *et al.*, 2007). We engineered a cell model to study the role of IFITMs in this type of cancer; SiHa cells originating from squamous cell carcinoma of cervix express IFITM1 and IFITM3 at detectable endogenous levels with a higher expression upon IFNγ stimulation (SI Appendix, Fig.1C). As IFITM2 protein was detected neither at endogenous level nor after IFNγ stimulation in the parental wt-SiHa cells (SI Appendix, Fig. S1D), we genetically engineered IFITM1 and IFITM3 knockout cell line (hereafter referred to as IFITM1/IFITM3 null) by using CRISPR/Cas9 technology (SI Appendix, Fig. S1B). The immune-related IFITM1/2/3 proteins have conserved amino acid sequences and form homo- and hetero-oligomers (Siegrist, Ebeling and Certa, 2011; Winkler *et al.*, 2019) suggesting that they cooperate together. Thus, to rule out any possible functional redundancy of IFITM1 and IFITM3, here, we use the IFITM1/IFITM3-knockout cells. Hence, the sgRNA was designed to introduce indels into the target IFITM sequences flanking the first exon for IFITM1 and IFITM3 genes to produce aberrant or truncated products (SI Appendix, Fig. S1A). After puromycin selection, individual clones were screened by immunoblotting (data not shown). A representative IFITM1/IFITM3 null clone showed IFITM1/3 protein expression diminished under the level of detection by Western blot (SI Appendix, Fig. S1C) and immunofluorescence (SI Appendix, Fig. S1E and F). More details on the design and screen of this cell model can be found in our previous publication (Gómez-Herranz *et al.*, 2019).

**Figure 1.**
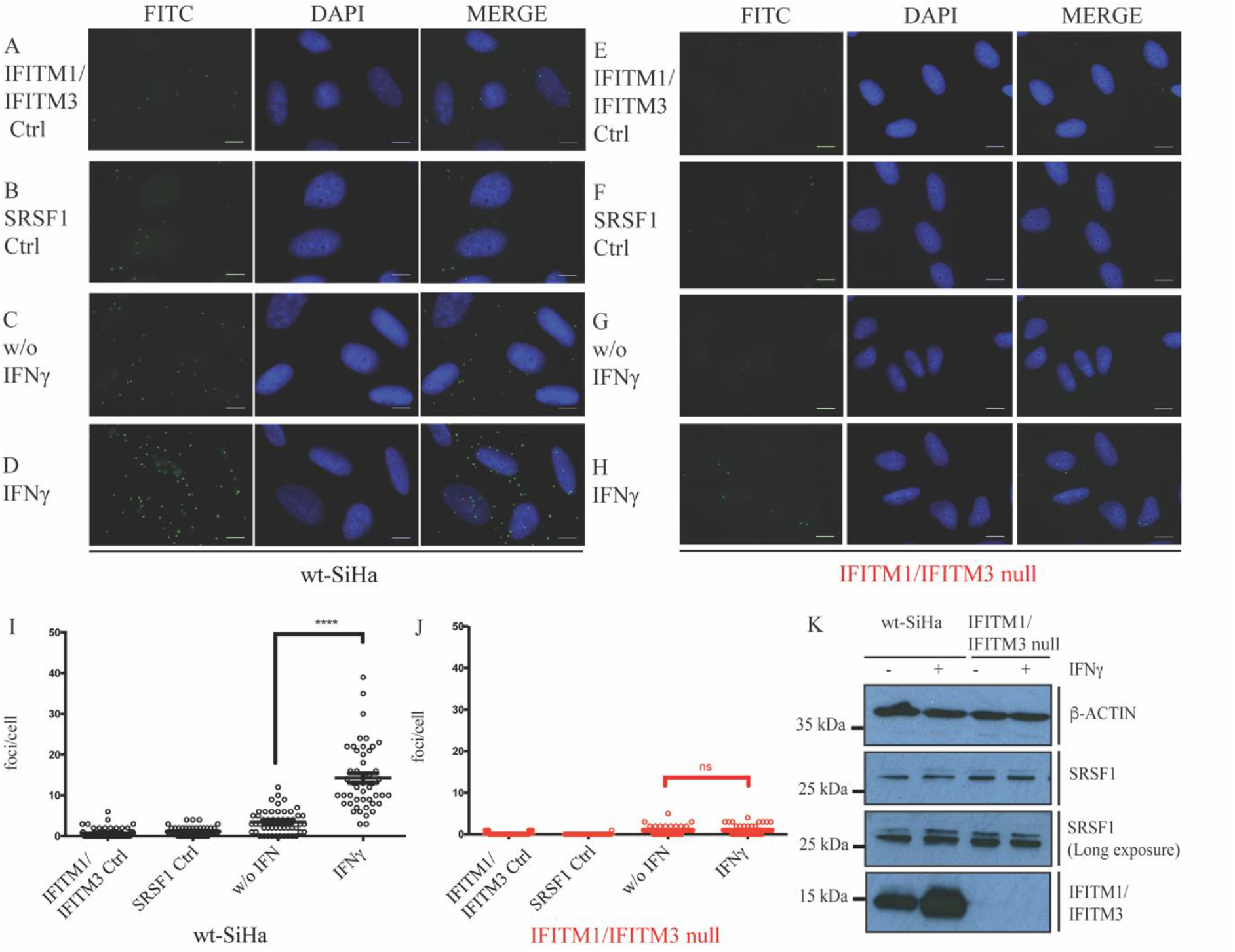
Evaluation of the IFITM1/IFITM3:SRSF1 protein-protein expression and interaction *in situ* after stimulation with IFNγ. Proximity ligation assays were used to study the endogenous interaction between SRSF1 and IFITM1/3 proteins in wt-SiHa (A-D) and IFITM1/IFITM3 null cells (E-H). FITC images identify the protein-protein association foci (depicted in green) and DAPI was used for nuclear staining (depicted in blue). (A-B and E-F) Cells were incubated as negative controls using IFITM1/IFITM3 or SRSF1 antibodies only. (C and G) Cells were incubated with both IFITM1/IFITM3 and SRSF1 antibodies to define protein-protein foci in non-stimulated cells. (D and H) Cells were incubated with both IFITM1 and SRSF1 antibodies to define protein-protein foci in IFNγ-stimulated cells. Representative quantification of the protein-protein interaction foci per cell in presence or absence of IFN stimulation in wt-SiHa cells (I) and IFITM1/IFITM3 null cells (J). At least 50 cells were counted in each condition. Statistical study was performed with one-way ANOVA and Bonferroni correction (****p < 0.0001; ns, not significant). n=3. Scale bar: 10 μm. (K) Immunoblots examining the endogenous levels of SRSF1 protein in non-stimulated or IFNγ-stimulated wt-SiHa cells or IFITM1/IFITM3 null cells.

### Interaction of IFITM1 protein with the cytoplasmic translation factor protein SRSF1

IFITM1 is the main IFITM family member reported in tumor progression and poor prognosis for many human cancer types (Györffy *et al.*, 2008b; Seyfried *et al.*, 2008; Weichselbaum *et al.*, 2008; Yu, Xie, Ng, Lum, M. Y. Cai, *et al.*, 2015; Borg *et al.*, 2016b; Ogony *et al.*, 2016; Sari, Y. G. Yang, *et al.*, 2016). Since IFITM interactome is not well defined, we asked whether new binding partners can emerge by employing Stable Isotope Labeling with Amino Acids in Cell Culture (SILAC) coupled to mass spectrometry (MS). The full-length IFITM1 gene was cloned into an SBP-tagged expression vector (Keefe *et al.*, 2001) to allow for its affinity capture from crude lysates after transfection (SI Appendix, Fig. S2A). The affinity-purification protocol using streptavidin-beads was designed to capture SBP-IFITM1 and associated proteins. Cells were isotopically labeled using SILAC RPMI media containing 13C labeled arginine and lysine amino acids (R6K6), while the SBP empty-vector used as a negative control was expressed in cells containing unlabeled amino acids.

Under these conditions, SBP-IFITM1 protein was detected in the affinity capture from cells grown in heavy media, at both 24 and 48 h after transfection (SI Appendix, Fig. S2B, lanes 4 and 8) and, more importantly, SBP-IFITM1 peptides were dominantly identified after MS analysis (SI Appendix, Fig. S2C and Table S1). This internal control highlights that the methodology is able to enrich and identify the IFITM1 bait protein.

We focused on the common interacting proteins detected in the SBP-IFITM1 purification between these two time points. The most common targets were the splicing regulatory factors within the SRSF superfamily of serine-arginine-rich splicing factors; SRSF1, SRSF2, SRSF3, SRSF6, and U2AF1 (SI Appendix, Fig. S2D; tryptic peptides are highlighted). Interestingly, one of the tryptic peptides derived from the SRSF1 isoform ASF-1, located in the alternatively spliced C-terminus characteristic for the cytosolic form that regulates protein translation on the ribosome (Maslon *et al.*, 2014) (SI Appendix, Fig. S2D; SRSF1 tryptic peptide is underlined in red).

SRSF1 is the archetype component of the SR family of splicing factors (Zheng *et al.*, 2020). Thus, the endogenous IFITM1 and SRSF1 co-association was further validated by proximity ligation assay (PLA), which can detect two endogenous protein-protein interactions in situ (Weibrecht *et al.*, 2010), not relying on the expression of exogenous vector constructs. In this case, IFITM1 and IFITM3 were simultaneously detected to capture all IFITM endogenous interaction with SRSF1. A basal level of IFITM1/IFITM3-SRSF1 foci can be observed in non-stimulated wt-SiHa cells (Fig 1C vs 1A or 1B; quantified in 1I). The stimulation with IFNγ for 24 h elevated the number of foci that predominate in the cytoplasm (Fig 1D vs 1C; quantified in 1I). The cytoplasmic localization of the majority of IFITM1/IFITM3:SRSF1 foci is consistent with the tryptic peptides for the cytoplasmic isoform SRSF1 detected in the affinity capture (SI Appendix, Fig. S2D; SRSF1 and Table S1).

As a control for specificity, the IFITM1/IFITM3 null cells did not exhibit IFITM1/IFITM3-SRSF1 foci, not even after IFNγ stimulation (Fig 1E-H; quantified in J). SRSF1 protein itself is not induced by IFNγ treatment (Fig 1K), nor is SRSF1 depleted in the IFITM1/IFITM3 null cells (Fig 1K). These data indicate that the IFITM1/3 proteins themselves are predominantly responsible for the IFNγ dependent induction of the SRSF1-IFITM1/IFITM3 foci (Fig 1D). The results suggest that the SBP-IFITM1 pull-down (SI Appendix, Fig. S2) captures authentically co-localizing IFITM1:SRSF1 complexes.

### Interaction of IFITM1/3 with HLA-B RNA

In our previous report, we described that endogenous HLA-B is deficiently expressed in IFNγ-stimulated cells lacking IFITM1/3 expression (Gómez-Herranz *et al.*, 2019). In addition, the proposed SRSF1 interacting protein (SI Appendix, Fig. S2) is involved in RNA binding and transport (39). Then, we asked whether IFITM1/3 associate with mRNA in vivo. With this purpose, we examined the interaction between IFITM1/3 with HLA-B RNA as a potential mechanism to account for IFITM1/3-dependent HLA-B protein synthesis in response to IFNγ. Protein-RNA proximity ligations (rISH-PLA) (Roussis, Myers and Scarlett, 2017) was used to measure IFITM1 protein-HLA-B RNA interactions. The IFITMs and HLA-B are ISG genes but the IFNγ-induced expression of HLA-B is impaired in the absence of IFITM1/3 expression (Gómez-Herranz *et al.*, 2019). Taking this into account, wt-SiHa cells and IFITM1/IFITM3 null cells were non-stimulated and IFNγ-stimulated for 24 h (Fig 2). Cells were processed as indicated in the methods and incubated with antibodies to IFITM1/3 and biotinylated probe designed to bind specifically the HLA-B mRNA.

**Figure 2.**
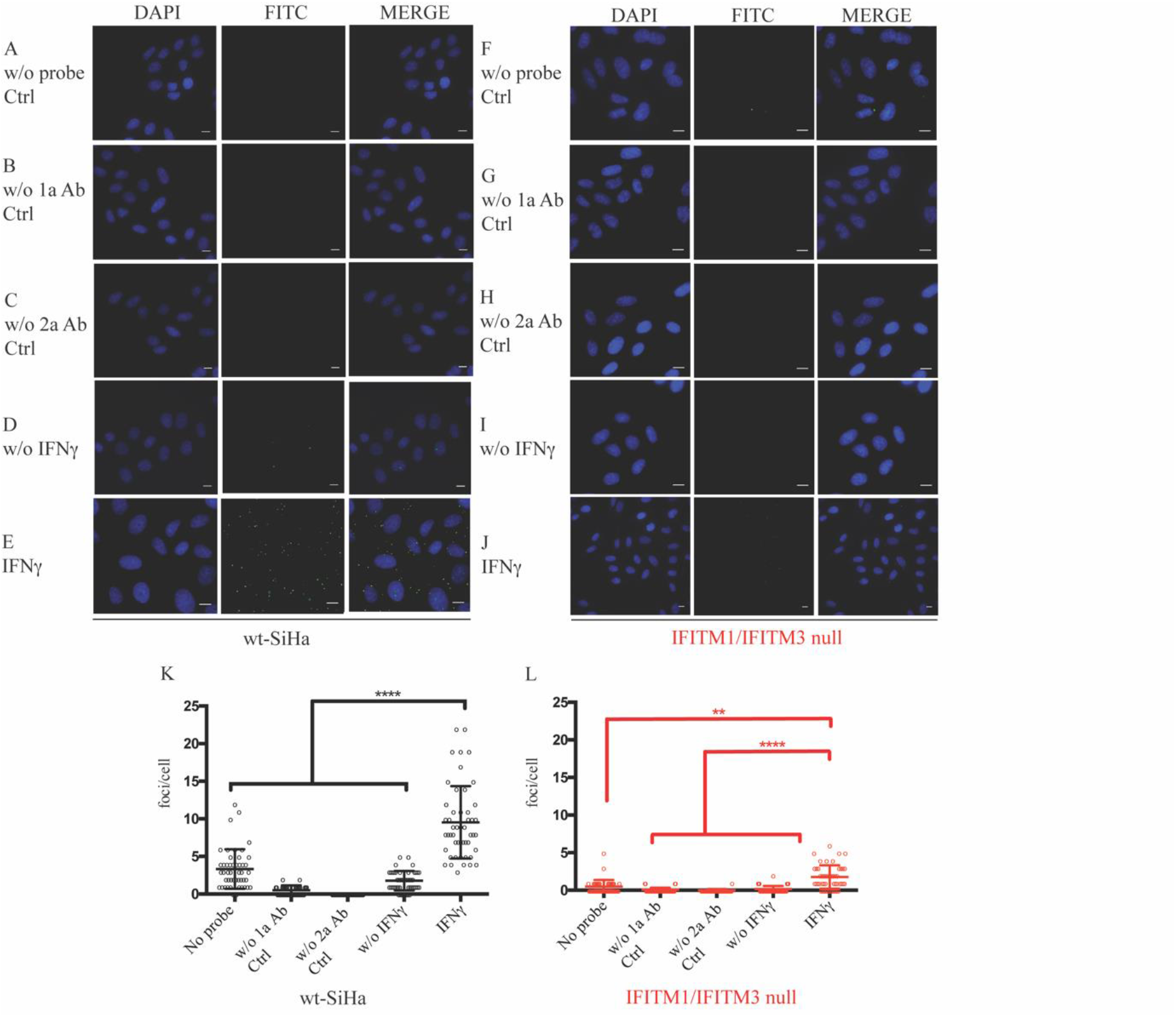
Evaluation of the IFITM1/IFITM3:HLA-B protein-RNA interaction *in situ* after stimulation with IFNγ by rISH-PLA. RNA in situ hybridization–PLA assays were used to study the endogenous interaction between HLA-B mRNA and IFITM1/3 proteins in wt-SiHa cells (A–E) and IFITM1/IFITM3 null cells (F-J). A biotinylated probe was designed to bind specifically the HLA-B mRNA. FITC images identify the protein–RNA association foci (green) and DAPI was used for nuclear staining (blue). (A–C and F-H) Cells were incubated as negative controls; without the RNA probe (A and F), using primary antibodies only (B and G), or secondary antibodies only (C and H). Cells were incubated with both IFITM1/IFITM3 and biotin antibodies to define protein–RNA foci in non-stimulated cells (D and I) or IFNγ-stimulated cells (E and J). Representative quantification of the protein–RNA interaction foci per cell in presence or absence of IFN stimulation in wt-SiHa cells (K) and IFITM1/IFITM3 null cells (L). At least 50 cells were counted in each condition. Statistical study was performed with one-way ANOVA and Bonferroni correction (****p < 0.0001, (**p < 0.01). n=3. Scale bar: 10 μm.

FITC images identify the protein–RNA association foci (green) and DAPI was used for nuclear staining (blue). Cells were incubated as negative controls (Fig 2A–C and F-H); without the RNA probe (Fig 2A and F), using primary antibodies only (Fig 2B and G), or secondary antibodies only (Fig 2C and H). Cells were incubated with both IFITM1/IFITM3 and biotin antibodies to define protein–RNA foci in non-stimulated cells (Fig 2D and I) or IFNγ-stimulated cells (Fig 2E and J). Endogenous interactions between HLA-B mRNA and IFITM1/3 proteins are shown in wt-SiHa cells (Fig 2A-E) and barely any interaction was observed in IFITM1/IFITM3 null cells (Fig 2F-J). Representative quantification of the protein–RNA interaction foci per cell in presence or absence of IFNγ stimulation in wt-SiHa cells (Fig 2K) and IFITM1/IFITM3 null cells (Fig 2L). Compiling both results (Fig 1 and Fig 2), the data suggest that IFNγ can induce the association of IFITM1/3 proteins with SRSF1 protein and HLA-B RNA. Bringing all together, these results start explaining why HLA-B protein synthesis is attenuated in the IFITM1/IFITM3 cells (Gómez-Herranz *et al.*, 2019).

### Transcriptomic analysis of the IFNγ response in wt-SiHA and an IFITM1/IFITM3 null cells

Next, we asked whether attenuated HLA-B protein synthesis in the IFNγ-stimulated IFITM1/IFITM3 null cells (Gómez-Herranz *et al.*, 2019) was related to reduced mRNA levels. This suggests that the dominant mechanism for IFITM1/3 protein signaling was transcription-dependent mechanisms. To address this question, a targeted, quantitative RT-PCR assay with biological triplicate samples from wt-SiHa and IFITM1/IFITM3 null cells non-stimulated or IFNγ-stimulated were measured, each in technical triplicate (SI Appendix, Fig. S3 I-IV-V). In addition to the HLA-B transcript, we also included additional IFNγ stimulated genes as controls; STAT1, B2M, and IRF1; further, we validate the IFITM1/3 knockout cells; IFITM1 and IFITM3. Of note, STAT1, B2M, IFITM1 and HLA-B are IRDS genes.

The positive controls highlight equivalent mRNA levels observed upon IFNγ induction of STAT1 (master signal transducer of the IFNγ pathway) and B2M (MHC class I subunit) in both wt-SiHa and IFITM1/IFITM3 null cells (Fig 3A I and II). This is consistent with our previous data, where we observed IFITM1/IFITM3-independent synthesis of STAT1 and B2M proteins using pulse-SILAC mass-spectrometry (Gómez-Herranz *et al.*, 2019). In addition, IRF1 (transcriptional activator of the secondary ISG response in the IFNγ pathway) was shown to partially attenuated in its synthesis in IFNγ-stimulated IFITM1/IFITM3 null cells using pulse-SILAC mass spectrometry (Gómez-Herranz *et al.*, 2019); its mRNA levels are also equivalently induced by IFNγ in both cell lines (Fig 3A V). Moreover, there is a reduction of both IFITM1 and IFITM3 mRNA abundance in IFNγ-stimulated IFITM1/IFITM3 null cells compared to wt-SiHa cells (Fig 3A III and 3 IV, respectively). These data might be expected if gene editing creates indels that result in stop codon mutations which cause mRNA degradation by NMD (Tuladhar *et al.*, 2019).

**Figure 3.**
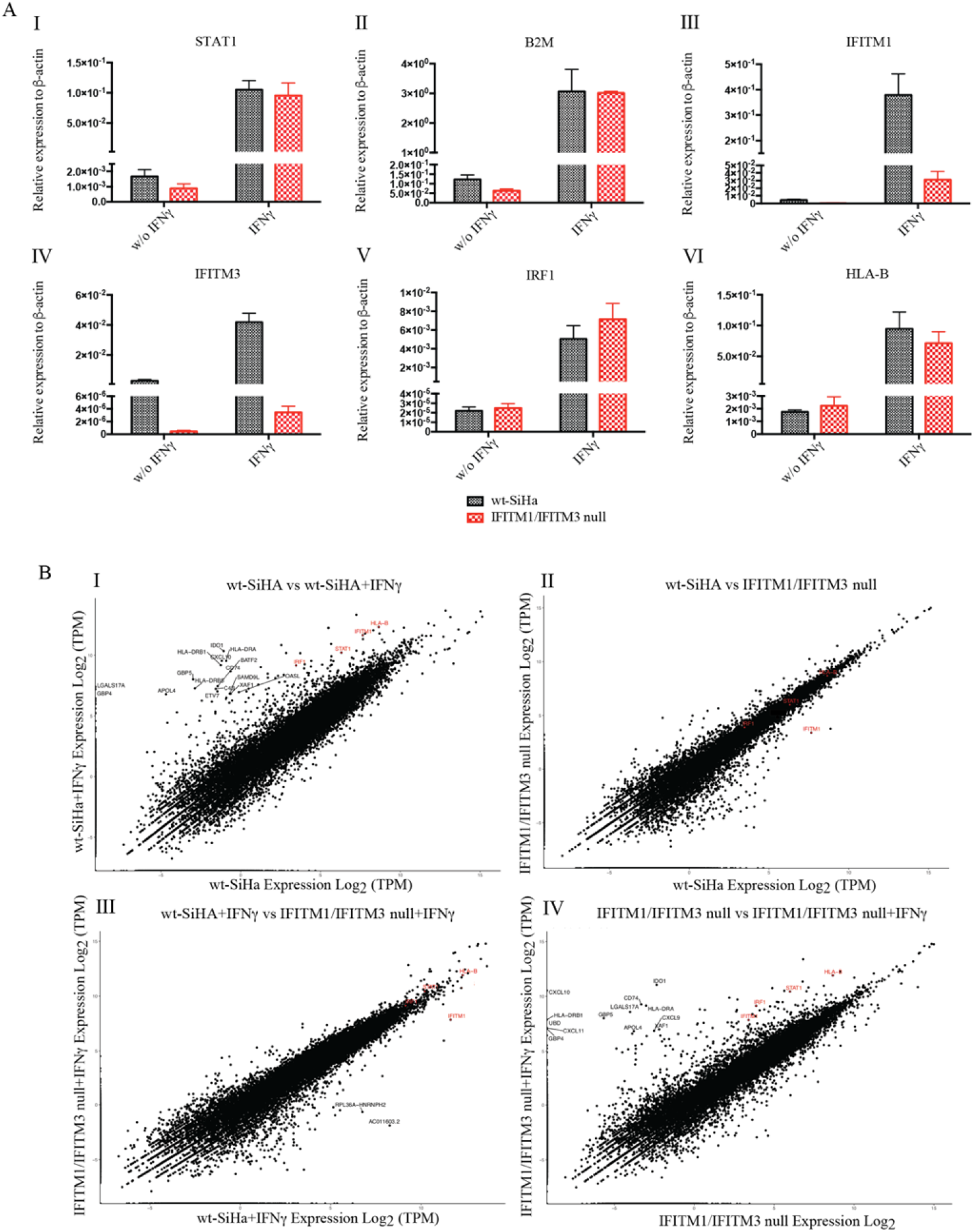
Validation of the transcript levels of IFNγ-stimulated genes in wt-SiHa cells and IFITM1/IFITM3 null cells unstimulated or IFNγ-stimulated for 24 h. STAT1 (I), B2M(II), IFITM1 (III), IFITM3 (IV), IRF1 (V), HLA-B (VI) mRNA quantification by RT-qPCR in wt-SiHa (in black) and *IFITM1/IFITM3* null cells (in red). Measurements were performed in untreated cells and after stimulation with 100 ng/ml IFNγ for 24 h. Error bars are a representation of the variability between three biological replicates. Each biological sample was run in three technical replicates. β-ACTIN was used to normalize the mRNA expression between samples. Statistical study was performed with two-way ANOVA and Bonferroni correction (****p < 0.0001; ***p < 0.001; ** p <0.01; ns, not significant). B. Comparison of total transcript count in wt-SiHa cells and IFITM1/IFITM3 null cells. Fastq files were imported into CLCBio Genomics workbench 12.0. The sequencing reads were used as the input file and RNA-seq analysis tool was used as the input for Create Fold Change Track drop down tab. All transcript reads detected were taken to generate the final transcript count for each gene (SI Appendix, Table S2). Comparisons of all transcripts identified are plotted for the following conditions: (I) non-treated wt-SiHa cells vs IFNγ-stimulated wt-SiHa cells, (II) non-treated wt-SiHa cells vs non-treated IFITM1/IFITM3 null cells, (III) IFNγ-stimulated wt-SiHa cells vs IFNγ-stimulated IFITM1/IFITM3 null cells, (IV) non-treated IFITM1/IFITM3 null cells vs IFNγ-stimulated IFITM1/IFITM3 null cells. Each plot has highlighted in red STAT1, IFITM1, IRF1, ISG15 and HLA-B transcripts. Other significantly induced transcripts are additionally highlighted in black. Transcript expression is measured in Log2 of transcripts per million (TPM).

Although HLA-B protein synthesis was not detected after IFNγ stimulation in an IFITM1/IFITM3 null cells using pulse-SILAC mass spectrometry (Gómez-Herranz *et al.*, 2019), there was an equivalent 100-250 fold increased relative levels of HLA-B mRNA in both the wt-SiHa cells and IFITM1/IFITM3 null cells (Fig 3A VI). This later result suggests that a transcription-independent mechanism is driving the reduced synthesis of HLA-B proteins after IFNγ stimulation in IFITM1/IFITM3 null cells. Since HLA-B mRNA are similarly induced by IFNγ in the IFITM1/IFITM3 null cells compared to wt-SiHA cells (Fig 3A VI), we next focused on understanding what might be linked to defects in protein synthesis in the IFITM1/IFITM3 null cells.

After these preliminary results, using the same RNA preparations used for RT-PCR analysis (Fig 3A), we performed global gene expression analysis by RNA-seq (Fig 3B) to investigate whether lack of IFITM1/3 expression, overall, changes other mRNA transcript products. Three biological replicates of wt-SiHa cells and the isogenic IFITM1/IFITM3 null cells were either left untreated (control) or IFNγ-stimulated for 24 h (SI Appendix, Fig. S3 I). Then, RNA from three biological replicates (non-stimulated or IFNγ-stimulated cells) was pooled, extracted, and processed by next generation RNA sequencing of polyA+ enriched RNA (SI Appendix, Fig. S3 II). The RNAseq data was annotated using CLC Genomics Workbench (12.0) (SI Appendix, Fig. S3 III and Table S2).

The wt-SiHa cells exhibited the expected induction of standard IFNγ-inducible genes, including HLA-B, IFITM1, STAT1, IRF1 (Fig 3B; gene names highlighted in red). Loss of IFITM1/3 protein expression does not decrease these transcripts (Fig 3B IV; highlighted in red). In addition to these classic IFNγ-inducible genes, other highly-induced transcripts after IFNγ treatment are also revealed to be IFITM1/IFITM3-independent; these include HLA-DRB1, HLA-DRA, CXCL10 and GBP5 (Fig 3 I and IV, highlighted in black). These genes are related to tumor immunity or pathogen restriction (Mach *et al.*, 1996; Soejima and Rollins, 2001; Feng *et al.*, 2017). Nonetheless, globally, no major differences were observed in the total amount of transcripts in the cells lacking of IFITM1/3 expression compared to wt-SiHa cells (Fig 3B).

By comparing the mRNA ratio in wt-SiHa cells to IFITM1/IFITM3 null cells, the dominant suppressed gene detected was IFITM1 (Fig 3B II and III; highlighted in red). This is consistent with; IFITM1 being a dominant interferon-inducible target and IFITM1 gene mutation through guide RNA editing (as described in (Gómez-Herranz *et al.*, 2019)) reduces the basal levels of the IFITM1 mRNA.

Furthermore, we focused on the mRNA abundance of the 31 genes composing the IRDS cluster. The HLA-B is an IRDS gene whose expression seems to be mediated by IFITM1/3; therefore, we investigated whether other transcripts originated from the IRDS genes may be affected. IRDS genes derived from Table S2 are a subset of IFN responsive genes linked to chemo-radiation resistance that include IFITM1 (but not IFITM3) and HLA-B (Erdal *et al.*, 2017). In this regard, a heat map quantifying the expression of the IRDS genes are highlighted in SI Appendix, Fig. S4). Overall, there is a substantial induction of IRDS genes after IFNγ stimulation in both, the wt-SiHa cells and IFITM1/IFITM3 null cells (SI Appendix, Fig. S4).

### Reduction in 80S ribosomal RNA levels in the IFITM1/IFITM3 null cells using sucrose gradient sedimentation

Inconsistent with the transcriptomic analysis results, we aimed to investigate whether IFITM1/3 proteins can modulate HLA-B expression by other mechanisms not involving cellular transcription. We previously have shown that HLA-B interacts with IFITM1/3 (Gómez-Herranz *et al.*, 2019) and, in this report, we have identified a new interacting partner for IFITM1, SRSF1, which interestingly mediates in the translation regulation (39). Taking all together, we investigate whether IFITM1/3 proteins may modulate HLA-B expression through intervention in the translational process.

In the attempt to study the relation of IFITM1/3 with HLA-B protein translation, we first asked whether IFITM1/3 proteins are present in the ribosome by studying their potential interaction with RPL7a protein, a ribosome-specific protein used as a subcellular marker. RPL7a is a component of the 60S ribosomal (Robledo *et al.*, 2008) that associates with the endoplasmic reticulum (Wu, Liu and Lin, 2007). By performing a PLA, there is a significantly increased protein–protein association of IFITM1/IFITM3:RPL7a after IFNγ-stimulation in the cytosol compared to non-stimulated wt-SiHa cells (Fig 4C vs D and I). In contrast, there is almost no detectable signal in IFNγ-stimulated IFITM1/IFITM3 null cells (Fig 4H and J). It is worth mentioning that IFITM1/IFITM3 proteins accumulate perinuclearly upon IFNγ stimulation (SI Appendix, Fig. S1E, arrowhead in merge composition with IFNγ-stimulated in wt-SiHa cells), which is consistent with a possible ribosomal localization after the activation of the IFNg signaling.

**Figure 4.**
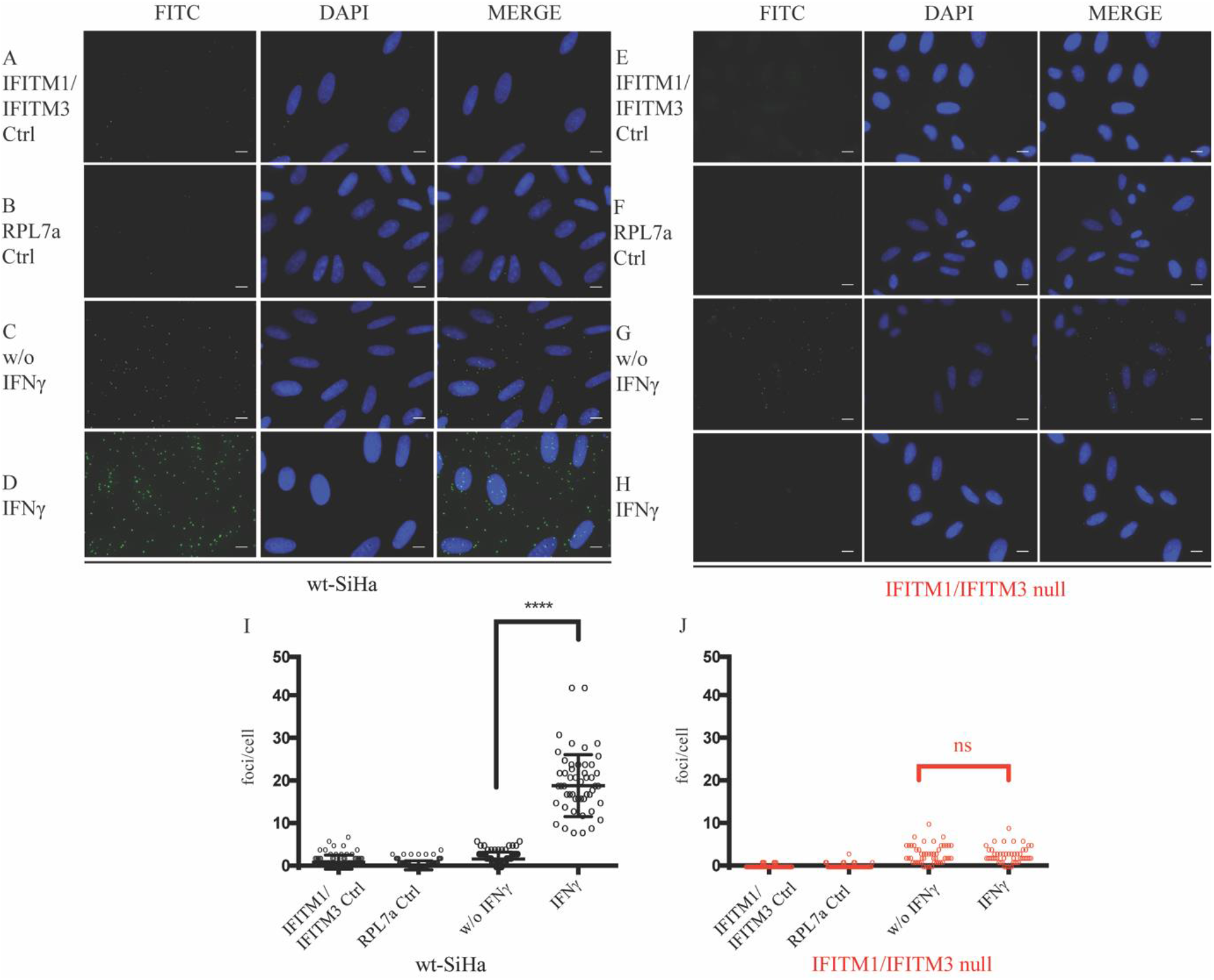
Evaluation of the IFITM1/IFITM3:RPL7a protein–protein expression and interaction in situ after stimulation with IFNγ. Proximity ligation assays were used to study the endogenous interaction between RPL7a and IFITM1/IFITM3 proteins in wt-SiHa (A–D) and IFITM1/IFITM3 null cells (E–H) FITC images identify the protein–protein association foci (green) and DAPI was used for nuclear staining (blue). (A–B and E–F) Cells were incubated as negative controls using IFITM1/IFITM3 or RPL7a antibodies only. (C and G) Cells were incubated with both IFITM1/IFITM3 and RPL7a antibodies to define protein–protein foci in non-stimulated cells. (D and H) Cells were incubated with both IFITM1 and RPL7a antibodies to define protein protein foci in IFN-stimulated cells. Representative quantification of the protein–protein interaction foci per cell in presence or absence of IFN stimulation in wt-SiHa (I) and IFITM1/IFITM3 null (J). At least 50 cells were counted for each condition. Statistical study was performed with one-way ANOVA and Bonferroni correction (****p < 0.0001; ns, not significant). n=3. Scale bar: 10 μm.

Given that IFITM1 can regulate HIV mRNA translation (Lee *et al.*, 2018), we set out to determine whether defects in ribosomal integrity was linked to defects in HLA-B protein synthesis (Gómez-Herranz *et al.*, 2019) in the IFITM1/IFITM3 null cells. An analysis of the ribosomal profile from the isogenic wt-SiHa and the IFITM1/IFITM3 null cell was first used to define ribosomal integrity. Then, cell lysates from IFNγ-stimulated cells were subjected to sucrose gradient sedimentation under conditions that maintain ribosome integrity (Fig 5). Immunoblots of the polysomal fractions indicated that IFITM1/3 proteins can be detected in the 40S, 60S, and 80S fractions from wt-SiHa cells but not from the IFITM1/IFITM3 null cells (Fig 5B).

**Figure 5.**
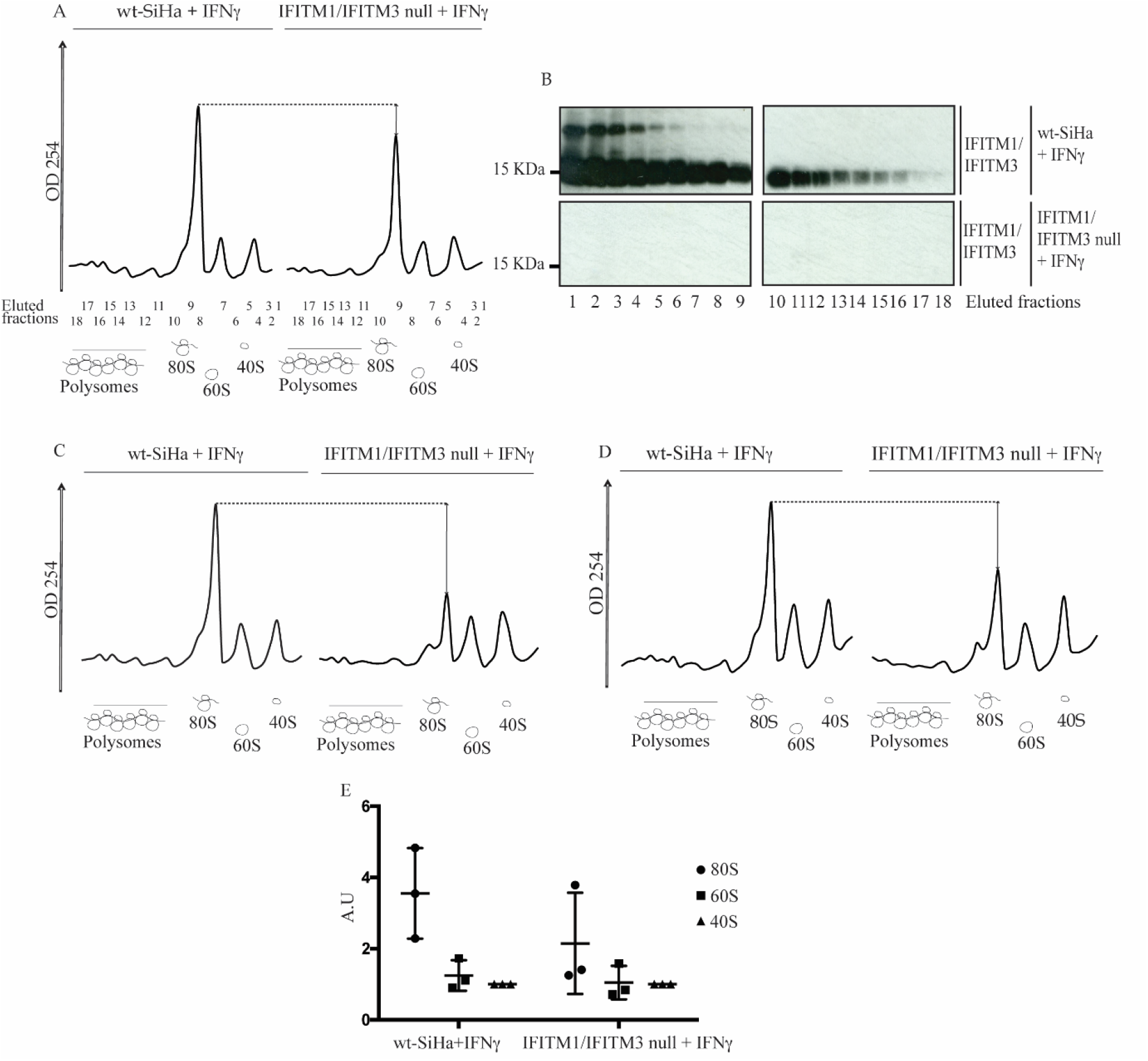
Ribosomal integrity analysis in IFNγ-stimulated wt-SiHa cells and IFITM1/IFITM3 null cells. (A, C, and D). The wt-SiHa cells and IFITM1/IFITM3 null cells were stimulated in three biological replicates with 100 ng/ml IFNγ for 24 h. The cells were lysed with ribosome stabilization buffer and applied to a 10-45% sucrose gradient to separate individual components of the large and small ribosomal subunits, and polysomal structures. Eluted fractions (numbered) were scanned at A254 nm and the diagram highlights the position of the 40S, 60S, 80S, and poly-ribosomal subunits. The dotted line highlights the reproducible reduction in the A254 signal in eluates from the IFITM1/IFITM3 null cells. (B) Following fractionation of material from (A), samples from fractions 1-18 were precipitated by trichloroacetic acid and they were analyzed for IFITM1/3 protein expression (enriched in all wt-SiHa fractions). (E) Quantitation of the ratio of 40S, 60S, and 80S A254 nm peaks in wt-SiHa cells and IFNγ-stimulated IFITM1/IFITM3 null cells.

The representative A254 nm peaks of 40S, 60S, 80S and polysomal fractions is depicted in Fig 5A. Interestingly, the A254 absorbance scan of the 80S subunit fraction in the IFITM1/IFITM3 null cells was reproducibly lower than in wt-SiHa cells in three independent biological replicates (Fig 5A, 5C, and 5D; quantified in 5E and SI Appendix, Fig. S5A-C). Remarkably, transient transfection of IFITM1 and IFITM3 genes for 24 h into the an IFITM1/IFITM3 null cells, in three independent biological replicates, can partially restore 80S ribosomal RNA levels as defined by increase in A254 nm (SI Appendix, Fig. S5D-F). These data suggest a selective impairment in the RNA component of the 80S ribosome in absence of IFITM1/3 expression. However, the defects observed in 80S integrity are not an irreversible feature of the IFITM1/IFITM3 null cell line.

## Discussion

Several studies have postulated that high expression of the IFN-stimulated IFITM1 and IFITM3 proteins can stimulate more aggressive growth of human cancers (Ogony *et al.*, 2016; Yang *et al.*, 2018; Lee *et al.*, 2020; Zhu *et al.*, 2020). In addition, IFITM1 is a component of the IRDS found up-regulated during the development of radiation resistance (Weichselbaum *et al.*, 2008). By contrast, IFITM1 activity is linked to growth suppression in cervical cancers (Yang *et al.*, 2007; Zheng *et al.*, 2017). In this regard, in our previous work, we identified a distinct subgroup of cervical cancer patients where IFITM1/IFITM3 proteins are highly expressed in cervical cancers. However, its expression is inversely correlated to the number of metastatic lymph nodes (Gómez-Herranz *et al.*, 2019). With this premise, we aimed to understand why patients who are IFITM1/IFITM3 double negative are also susceptible to developing aggressive tumors.

To broaden the limited molecular understanding of IFITM1 and IFITM3 function(s), in our previous report, we developed a MS assay to determine whether IFITM1 and IFITM3 play a role in the response of cells to IFN-induced protein synthesis (Gómez-Herranz *et al.*, 2019). As a result, we identified HLA-B, an MHC class I molecule, as one of the few dominant downstream effectors of IFITM1/3 (Gómez-Herranz *et al.*, 2019). Moreover, a decreased expression of HLA-B on the cell surface in cells with complete loss of IFITM1/3 expression was observed. Taking all together, the data suggested that loss of IFITM1/3 expression would attenuate antigen presentation *in vivo* (Gómez-Herranz *et al.*, 2019). However, in this current study, we are interested in which cellular process explains why HLA-B exhibits lowered synthesis in the IFITM1/IFITM3 null cells. With this, we could further comprehend other mechanisms by which the antigen presentation pathway is regulated.

To begin with, we start tackling this question by identifying new interacting partners for IFITM that could shed some light on which novel role IFITM1/3 proteins can be implicated. We identified multiple members of the SR family of splicing factors (SRSF1, SRSF2, SRSF3, SRSF6 and U2AF1) as novel candidates to associate with IFITM1 protein by employing affinity capture assay coupled to pulse-SILAC mass spectrometry (SI Appendix, Fig. S2). A Proximity ligation assay confirmed the association between IFITM1/3 and SRSF1 (Fig 1). However, immunoblot did not show major difference in the expression of SRSF1 in cells lacking of IFITM1/3 expression (Fig 1K). Interestingly, in addition to be implicated with the splicing machinery, the 5 splicing factors identified are implicated in the mRNA transport (Das and Krainer, 2014). SRSF2 is considered as nonnuclear to cytoplasmic shuttled protein, however, it has its C-terminal enriched in arginine and serine; these amino acid motifs are crucial for the nuclear exit of the mentioned slicing factors (SI Appendix, Fig. S2D; SRSF2). Thus, we study the possible implication of IFITM1/3 proteins regulating the transcript products. To address this possibility, the performed RNA expression analysis did not reveal a dominating defect in the HLA-B gene expression neither in the global transcript production by comparing IFITM1/IFITM3 null cell with wt-SiHa cells (Fig 3 and SI Appendix, Fig. S3 and Fig. S4). These data suggested that defects in HLA-B protein synthesis in cells lacking IFITM1/3 expression are not selectively due to suppression of IFN-stimulated gene induction even though IFITM1/3 associates with the HLA-B mRNA (Fig 2).

SRSF factors are implicating in guiding the mRNA products to the ribosome (Das and Krainer, 2014) and a ribosomal localization of IFITM1/3 proteins is also inferred due to the co-association of RPL7a and IFITM1/3 proteins using proximity ligation assays (Fig 4). RPL7a is a component of the 60S ribosomal subunit that contains nucleic acid-binding domains, and it is implicated in regulating the expression of mRNAs (Neumann *et al.*, 1995; Neumann and Krawinkel, 1997).

Next, we interrogated whether IFITM1/3 could be mediating selectively the protein synthesis, more representative for HLA-B, by modulating their translation. To find some answers, we evaluated the ribosomal profile using ultracentrifugation sedimentation of cell lysates from IFNγ-stimulated wt-SiHa cells and IFITM1/IFITM3 null cells to isolate ribosomal constituents. Our results suggest a specific role for IFITM1/IFITM3 in regulating the integrity of 80S ribosomal subunit, restored with the exogenous transient expression of IFITM1/3 (Fig 5 and SI Appendix, Fig. S5). During the course of our studies, it was shown that IFITM protein expression reduces HIV-1 viral protein synthesis by preferentially excluding viral mRNA transcripts from translation (Lee *et al.*, 2018). This study supports our vision where a new exciting role for IFITM family members operating at the level of protein translation is emerging. Although we focused on the role of IFITM1/3 regulating HLA-B, there may be other cancer and virus originated proteins whose expression may be mediated by IFITM1/3.

Here, we are starting to reveal an alternative novel role for IFITM1/3 proteins that may help to understand the complicated mechanism by which MHC class I molecules are regulated. Further direction will address the question on how specifically IFITM1/3 mediates the HLA-B expression; whether is required to form the protein complexes guiding mRNA, guides the mRNA to the ribosome, excludes the mRNA loading into the ribosome or a combination of all.

However, our findings emphasize the importance of IFITM1/IFITM3 to mediate HLA-B protein expression through interactions with its mRNA. Such RNA-dependent effects on HLA-B synthesis might impact on the global number of neo-antigens presented to CD8+ T cells, otherwise promoting immune scape. From a cervical cancer perspective, MHC class I depletion is not correlated with HPV positive tissues (Connor and Stern, 1990; Cromme *et al.*, 1993); it is suggested that these events are originated at different stages of cervical cancer progression. Independent studies have identified impaired MHC class I expression in cervical tumors (and other cancer types); particularly loss of MHC class I molecules occur in metastatic lesions compared to primary tumors (Torres *et al.*, 1993; Honma *et al.*, 1994), including chromosomal aberrations (Koopman *et al.*, 2000). This is of particular relevance for our study as it supports the hypothesis whereby IFITM1/3 are able to regulate, at least partially, the expression of MHC class I proteins. Remarkably, several studies revealed HLA expression deficiencies in lymph node metastases compared to primary tumor (Cromme *et al.*, 1994; Ferns *et al.*, 2016) which is consistent with the concept that lack of IFITM1/IFITM3 would lead to diminished HLA expression (Gómez-Herranz *et al.*, 2019).

## Supporting Information

This article contains SI Appendix; Fig. S1: Workflow for generating IFITM1/3-knockout SiHa cells using CRISPR/Cas9 gene editing technology, Fig. S2: Identification of IFITM1 interacting proteins, Fig S3: Workflow followed to analyze the transcriptome in wt-SiHa cells and IFITM1/IFITM3 null cells, Fig. S4: Heat map representation of the mRNA induction of the 31 IRDS genes, Fig. S5: Original scans for the mRNA trace after sucrose density gradient fractionation in six independent replicates, Table S1: List of proteins enriched in SBP-IFITM1 pull down, Table S2: Generation of RNA seq datasets from the indicated cell lines using CLCBio Genomics workbench 12.0.

## Acknowledgments

We would like to thank all past and current member of the laboratory for their contribution to scientific discussions and giving us so much support throughout all these years of research. A special thank you to Dr. Marta Nekulova for engineering the CRIPSR/Cas9 IFITM1/IFITM3 null cells and to Dr. Magadalena M. Maslon and Cáceres group for the help provided during polysome fractionation.

## SI APPENDIX

### Supplementary Information

**Figure S1.**
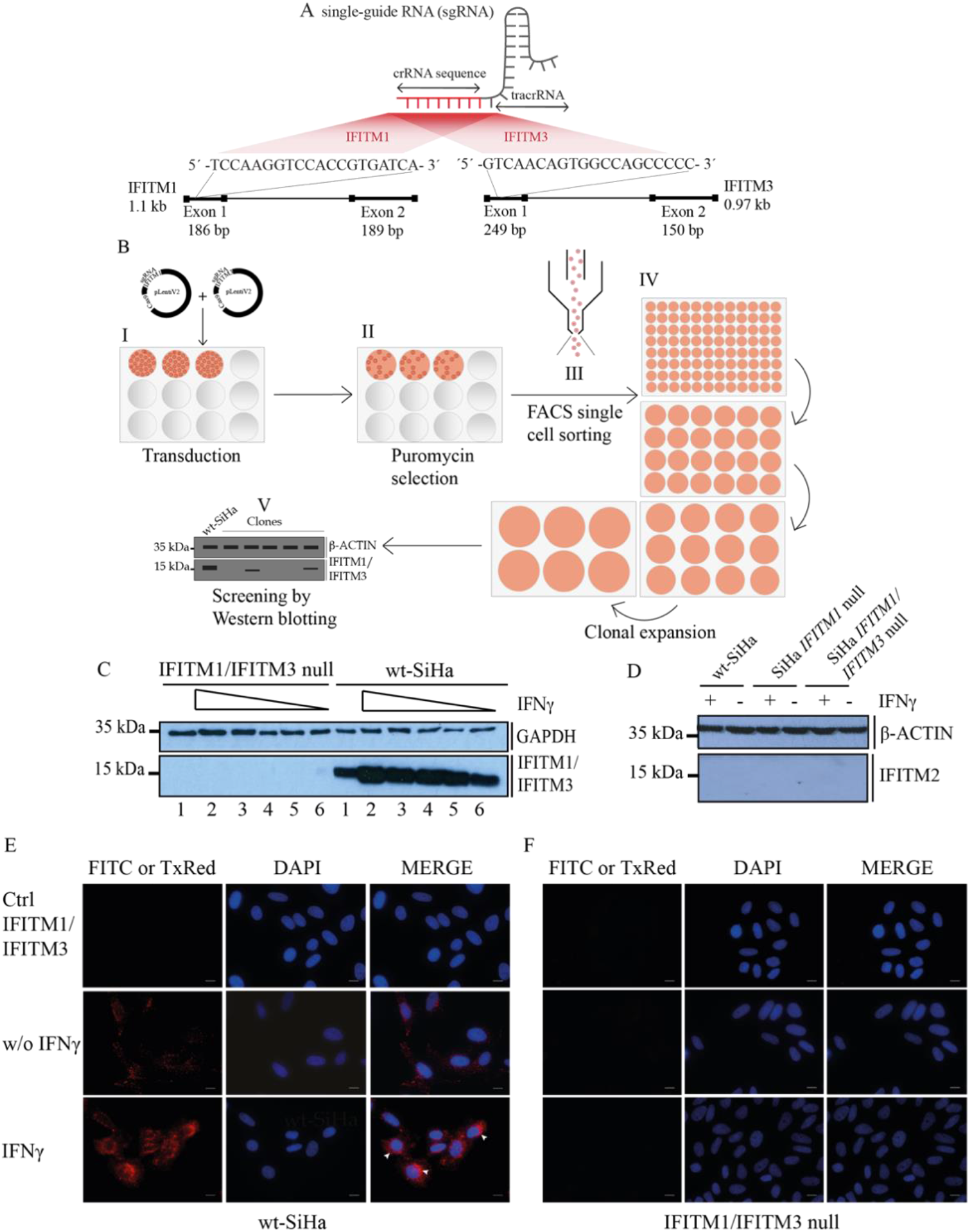
Workflow for generating IFITM1/3-knockout SiHa cells using CRISPR/Cas9 gene editing technology. (A) Schematic of the CRISPR/Cas9 system. The sgRNA sequence was designed to target the genes of interest within the exon 1; IFITM1 (5’-TCCAAGGTCCACCGTGATCA-3’) and IFITM3 (5’-GTCAACAGTGGCCAGCCCCC-3’). (B) The wt-SiHa cells were transduced with pLentiV2 plasmid containing the Cas9 cassette, IFITM1 sgRNA or IFITM3 sgRNA to generate IFITM1/IFITM3-knockout cells (I). The IFITM1/IFITM3 null cells were generated by transducing pLentiV2 plasmid containing Cas9 cassette and IFITM3 sgRNA into IFITM1 null cells. The next day, medium with lentivirus was removed and the cells were resuspended in fresh media containing 10 μg/ml puromycin (II). Puromycin selection was continued for about two weeks, after which single cells were sorted by FACS into 96-well plates (III). These cells were propagated to form colonies originating from a single cell (IV) and cell clones were tested by Western blotting (V). (C) Dose dependent titration in wt-SiHa and IFITM1/IFITM3 null cells with IFNγ-stimulation for 24 h: non-treated (lane 1), 2000 (lane 2), 1000 (lane 3), 100 (lane 4), 50 (lane 5), and 10 (lane 6) ng*/*ml. Cells were tested by Western blotting using the IFITM1/IFITM3 antibody for detection (approximately 15 kDa). GAPDH (approximately 35 kDa) was used as a loading control. (D) IFITM2 induction was tested by comparing endogenous protein expression to cells stimulated with 100 ng/ml IFNγ for 24 h. IFITM2 protein expression was not detected despite IFNγ-stimulation. Cells were tested by Western blotting using the IFITM2 antibody for detection (approximately 15 kDa). β-ACTIN (approximately 35 kDa) was used as a loading control. Wt-SiHa cells (E) and IFITM1/IFITM3 null (F) were grown to 80% confluency and fixed with 4% (w/v) paraformaldehyde, permeabilised using 0.25% Triton X-100 and blocked with 3% (w/v) BSA. Immunofluorescence was performed detecting IFITM1/IFITM3 protein in non-stimulated cells (w/o IFNγ) or cells stimulated with 100 ng/ml IFNγ (IFNγ) for 24 h. The negative control (Ctrl IFITM1/3) was performed by solely staining with anti-mouse Alexa Fluor 594 secondary antibody. Arrowhead coloured in white (D; merged column, IFNγ row) indicate the perinuclear distribution of IFITM1/IFITM3 protein in wt-SiHa upon IFNγ stimulation. Scale bar: 10 μm.

**Figure S2.**
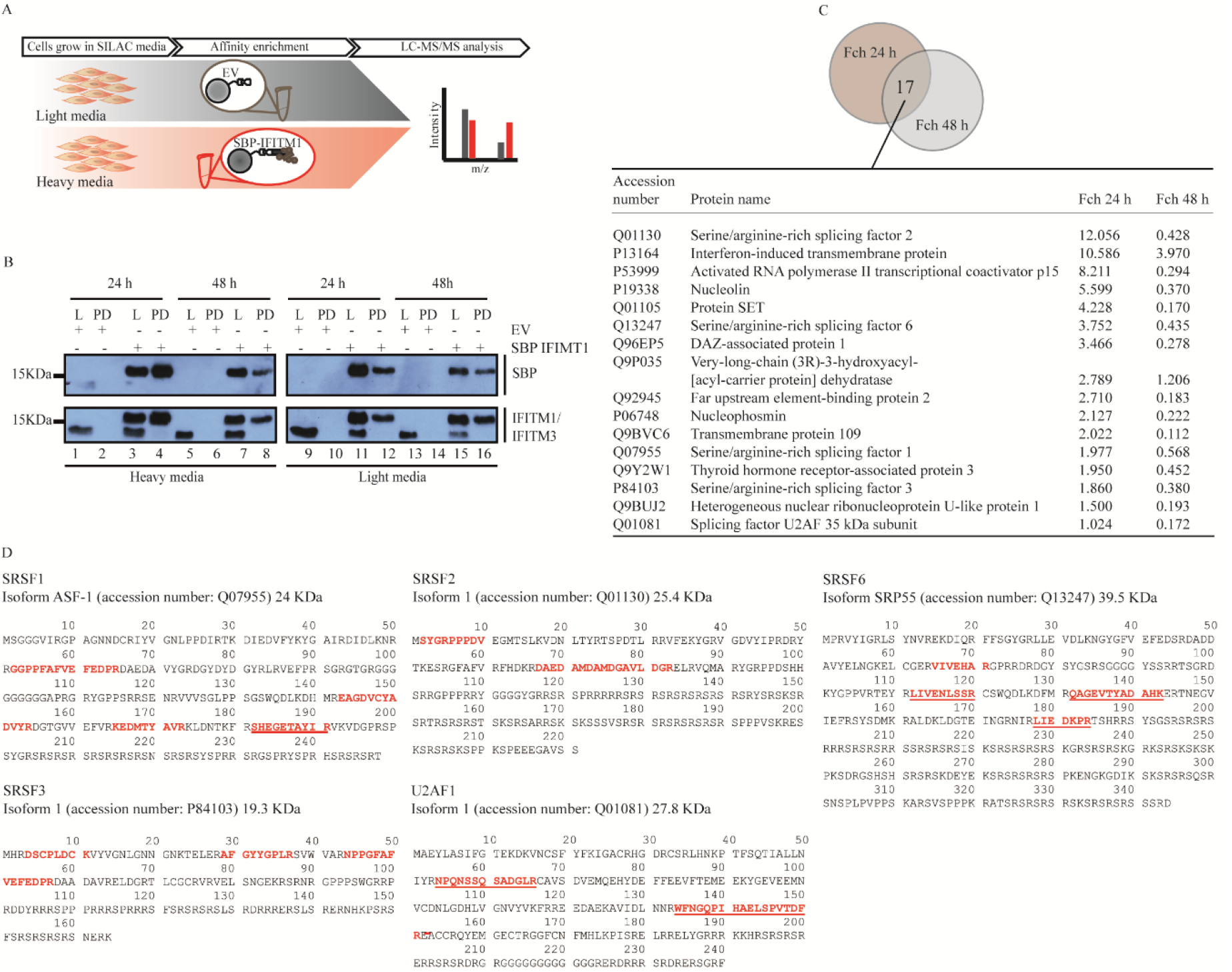
Identification of IFITM1 interacting proteins. (A) SILAC workflow describing the stages followed for identifying the IFITM1 binding proteins using SBP-tagged IFITM1 and SBP empty-vector control; stage include cell labelling, transfection, affinity purification using streptavidin beads, elution of the IFITM1 complexes, FASP and LC-MS/MS analysis. (B) An immunoblot of lysates from cells transfected with SBP-tagged IFITM1 or SBP empty vector. Cells were harvested after 24 and 48 h following SBP-IFITM1 or SBP-vector transfection. Transfections were performed in cells grown with heavy and light SILAC media. Lanes were loaded with Lysate [L] and Pull-down [PD]. SBP-tagged IFITM1 was detected in lysates of transfected cells grown in heavy (R6K6) media (lanes 1-8) or light (R0K0) media (lanes 9-16). The transfected SBP-IFITM1 protein was detected in the lysate using either anti-SBP antibody (top panel) or IFITM1/IFITM3-specific monoclonal antibody (lower panel; lanes 3, 7, 11 and 15). SBP-IFITM1 protein was detected by a migration at 19 kDa and the endogenous IFITM1/IFITM3 proteins migrated at a position of 13.9 kDa and 14.6 kDa, respectively. The transfected SBP-IFITM1 protein was detected in the lysate at either 24 or 48 h post-transfection into cells with heavy media (lanes 3 and 7) or light media (lanes 11 and 15). (C) Overlapping of proteins detected by MS from 24 or 48 h post transfection and affinity purification of SBP-IFITM1 including the top proteins enriched in the IFITM1 enrichment (from SI Appendix, Table S1). (D) Identified SRSF peptides in SBP-IFITM1 protein enrichment after affinity purification in isotopically labelled wt-SiHa cells. Heavy isotopically labelled tryptic peptides identified from the SBP-IFITM1 affinity enrichment are highlighted in red for the SRSF family of proteins (SRSF1, SRSF2, SRSF3, SRSF6 and U2AF1). Underlined unique peptides confirm that these proteins are SRSF isoforms reported to shuttle between the cytoplasm and nucleus.

**Figure S3.**
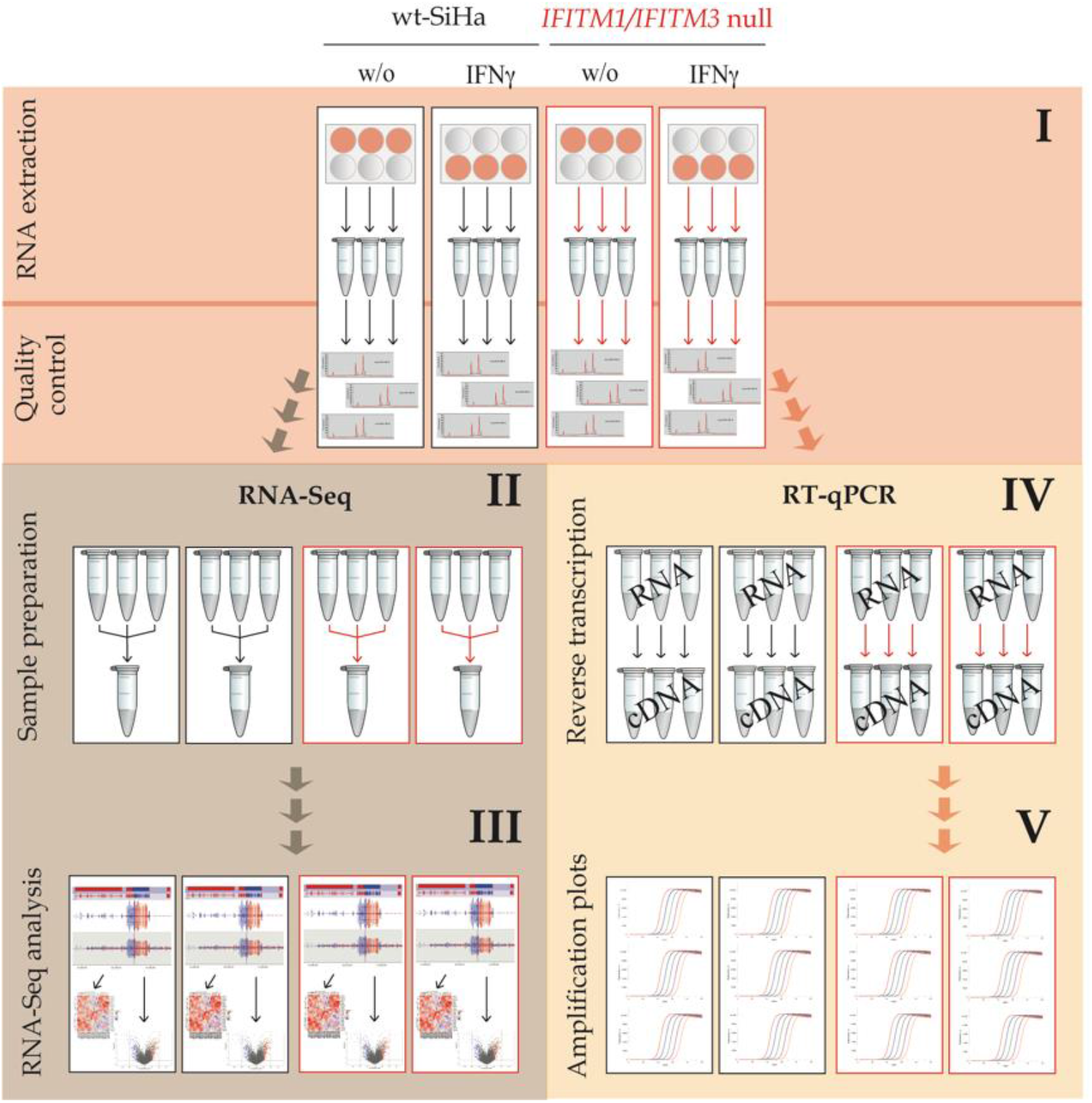
Workflow followed to analyze the transcriptome in wt-SiHa cells and IFITM1/IFITM3 null cells. (I) RNA was extracted from non-stimulated or IFNγ-stimulated cells using three biological replicates. Aliquots of total RNA from the three biological replicates were used for (II) next-generation RNA Seq analysis using the Illumina PE100-125 and HiSeq2500 sequencing platform. PolyA+ RNA pools were purified and used to generate cDNA for fragmentation, PCR amplification, and DNA sequencing according to Illumina methodologies. (III) The fastq files representing 20 M reads were imported and analyzed using CLC Genomics Workbench with GRCh38 as a reference genome. The global analysis of IFNγ responsive gene sets is plotted in Fig 3B I-IV and as a subset of the IFNγ responsive proteins, IRDS, as a heat map (SI Appendix, Fig. S4). (IV) Remained material from the RNA extraction was further used to analyze transcripts of interest by RT-qPCR using the three biological replicates (Fig 3). For this purpose, the RNA was reverse transcribed into cDNA and (V) sequentially quantified and compared between samples.

**Figure S4.**
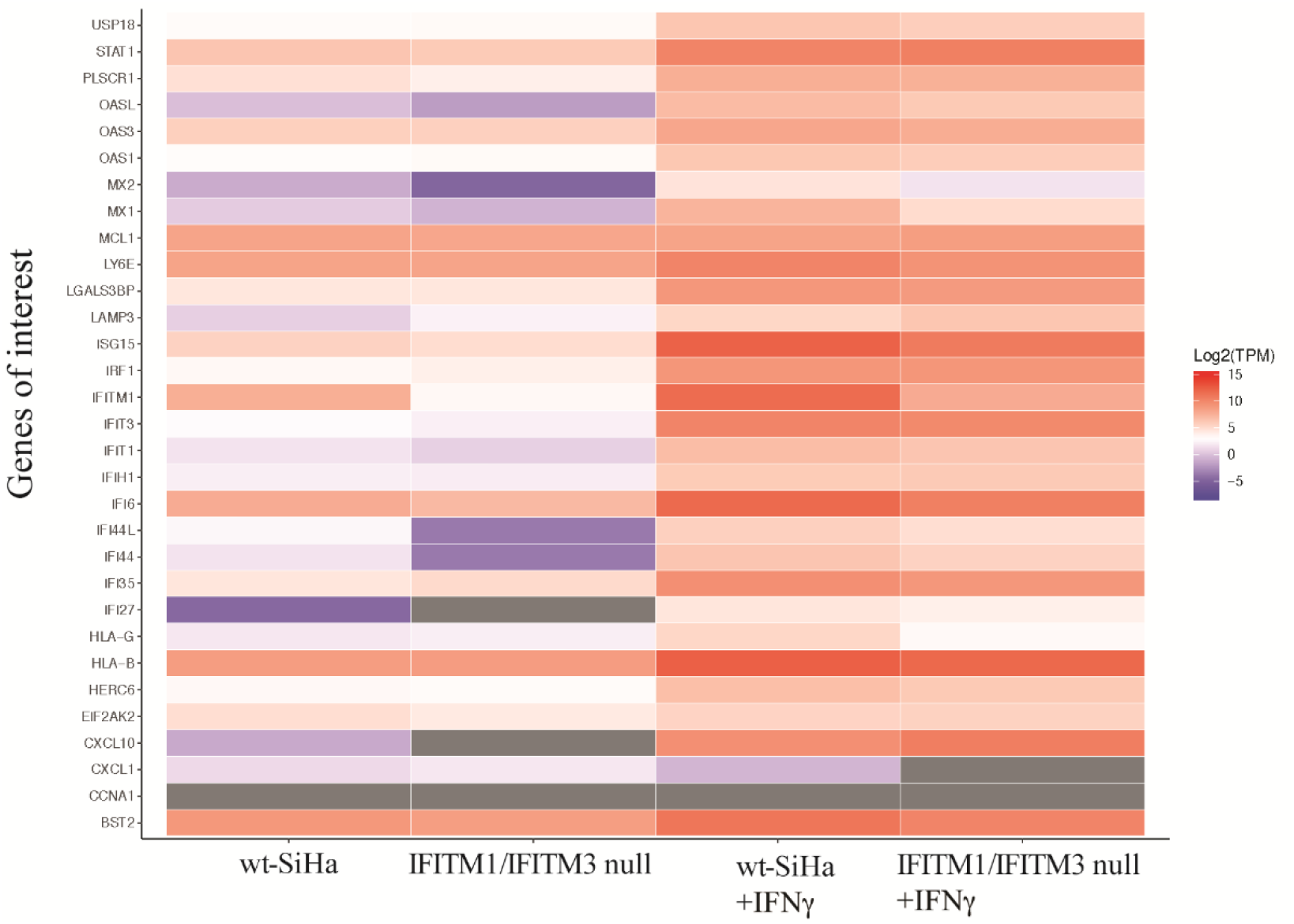
Heat map representation of the mRNA induction of the 31 IRDS genes. The x-axis contains the wt-SiHa and IFITM1/IFITM3 null cells non-stimulated or IFNγ-stimulated with 100 ng/ml for 24 h. The y-axis contains transcript expression corresponding to the 31 IRDS genes. Color scale units are log2 (TMP), becoming red when it is highly expressed and purple to blue for non-expressed and under-expressed values. Values in grey color correspond to genes which TPM was equal to 0. TPM=transcript per million. The heatmap was developed by taking the log2 (TPM) count of the 31 IRDS genes; representation of the genes of interest (IRDS) is extracted from SI Appendix, Table S2. R version 3.5.3 and ggplot2 3.2.1 was to create the heatmaps.

**Figure S5.**
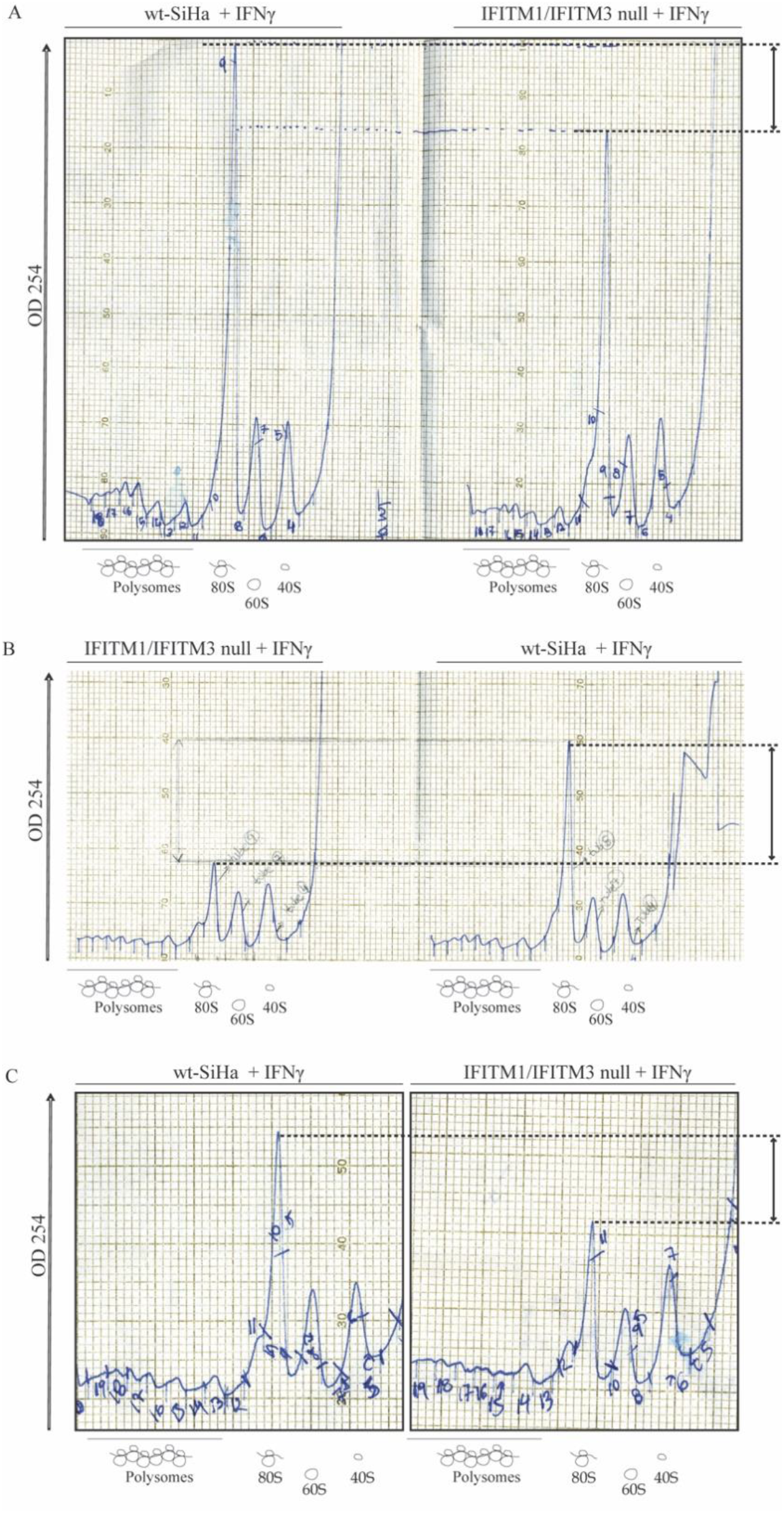
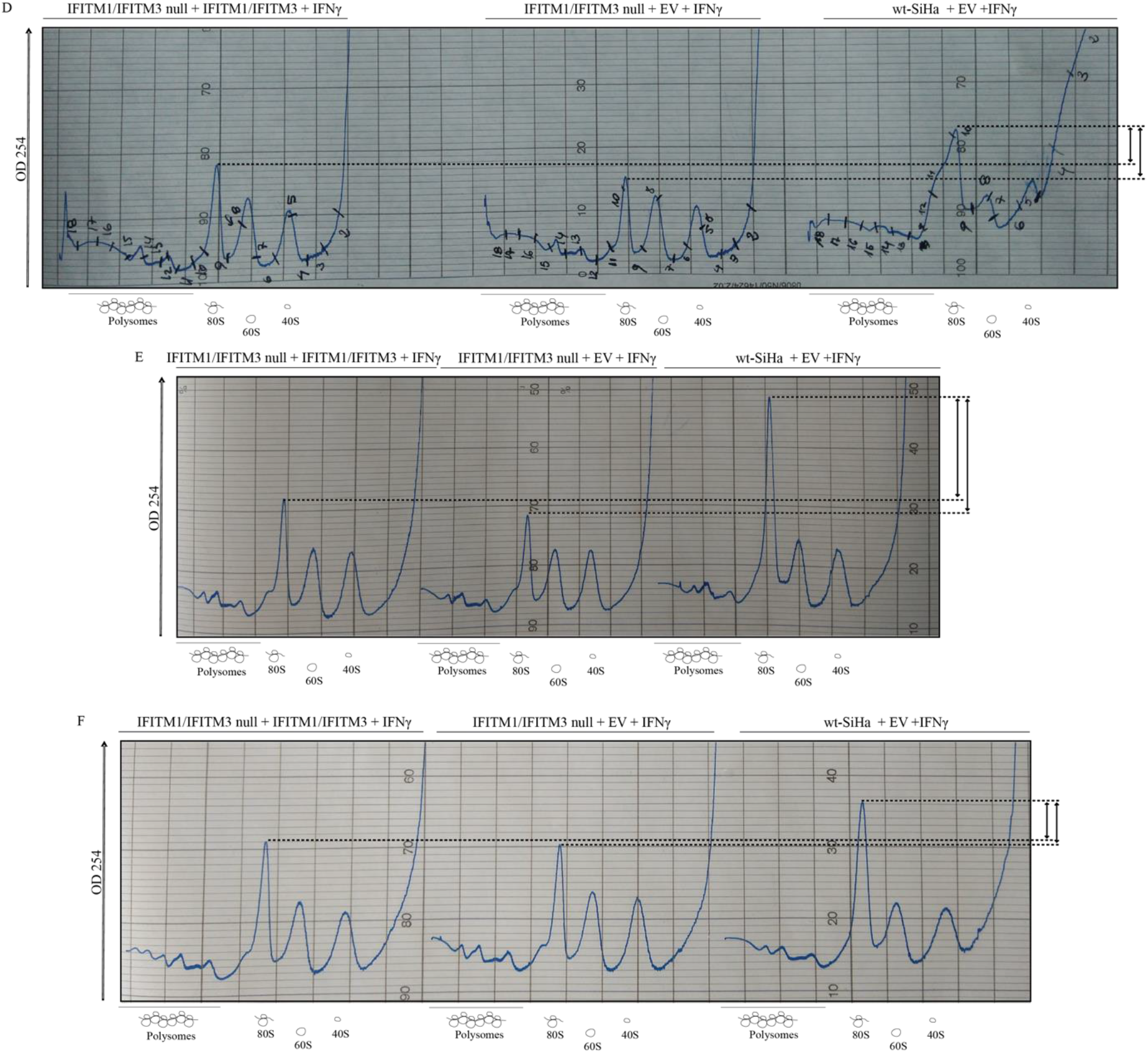
Original scans for the mRNA trace after sucrose density gradient fractionation in six independent replicates. (A–C) Sucrose density gradient was performed in wt-SiHa cells and IFITM1/IFITM3 null cells treated with 100 ng/ml IFNγ for 24 h to activate the IFN-protein synthesis response. A higher 80S peak (A254) was observed in wt-SiHa cells compared to in IFITM1/IFITM3 null cells in all three replicates (indicated with the dashed lines). (D-F) Sucrose density gradient was performed in wt-SiHa cells and IFITM1/IFITM3 null cells treated with 100 ng/ml IFNγ for 24 h. Cells were also transfected with IFITM1 and IFITM3, or the respective empty vector (EV) for 48 h to recover the lower 80S peak observed in the IFITM1/IFITM3 null cells (indicated with the dashed lines).

**Table S1. List of proteins enriched in SBP-IFITM1 pull down**. The precipitates at both 24 and 48 h transfection times were processed as in the experimental methods. The data in the multiconsensus report are represented, as in column: A. *Accession number,* B. *Description* (gene name). Summary from all analyzed samples. C. ΣCoverage (the number of amino acids in a protein sequence that were found in identified peptides), D. *Σ# Proteins* (number of proteins identified in the protein group; introduced is the master protein), E. *Σ# Unique Peptides* (number of peptides that are unique to a protein group), F. *Σ# Peptides* (the total number of distinct peptides in protein group), G. *Σ# PSMs* (the number of peptide spectrum matches, the total number of spectra used for the identification of the peptides belongs to the protein). 24 h time point; H. XCorr (the goodness of fit of experimental peptide fragments to theoretical spectra created from the sequence *b* and *y* ions)); I *Coverage,* J. # Peptides, K. *#PSM,* and L. Area (under the peak, value used for quantification). 48 h time point; M, XCorr; N *Coverage,* O # Peptides, P. *#PSM,* and Q. Area. The ratio of the relative peak intensities of the heavy to light peptides are highlighted at 24 h (R) and 48 h (U).

**Table S2. Generation of RNA seq datasets from the indicated cell lines using CLCBio Genomics workbench 12.0.** The fastq sequencing reads were used as the input file and RNA-seq analysis tool was used in the CLCbio Genomics workbench 12.0. All transcript reads detected were taken to generate the final transcript count for each gene. Comparisons of all transcripts were performed for the following cells: non-treated wt-SiHa cells (index 13 fastq files), IFNγ-stimulated wt-SiHa cells (index 14 fastq files), non-treated IFITM1/IFITM3 null cells (index 23 fastq files), and vs IFNγ-stimulated IFITM1/IFITM3 null cells (index 25 fastq files). The output excel file from the software in the columns represent: gene name; chromosome location; region ; identifier; wild type SiHa total counts; wild type SiHa RPKM; wild type SiHa TPM; wild type SiHa CPM; wild type SiHa IFNγ total counts; wild type SiHa IFNγ RPKM; wild type SiHa IFNγ TPM; wild type SiHa IFNγ CPM; IFITM1/IFITM3 null total counts; IFITM1/IFITM3 null RPKM; IFITM1/IFITM3 null TPM; IFITM1/IFITM3 null CPM; IFITM1/IFITM3 null IFNγ total counts; IFITM1/IFITM3 null IFNγ RPKM; IFITM1/IFITM3 null IFNγ TPM; IFITM1/IFITM3 null IFNγ CPM.

